# Advanced paternal age diversifies individual trajectories of vocalization patterns in neonatal mice

**DOI:** 10.1101/738781

**Authors:** Lingling Mai, Hitoshi Inada, Ryuichi Kimura, Kouta Kanno, Takeru Matsuda, Ryosuke O. Tachibana, Valter Tucci, Fumiyasu Komaki, Noboru Hiroi, Noriko Osumi

## Abstract

Infant crying is an innate communicative behavior that is frequently impaired in certain neurodevelopmental disorders (NDDs). Since advanced paternal age is a reported risk factor for NDDs in offspring, we evaluated the impact of a father’s age on early vocal development in C57BL/6J mice. We recorded and applied a unique combination of computational analyses to ultrasonic vocalizations (USVs) emitted by mouse pups sired by young and aged fathers. Our data showed that advanced paternal age reduced the number and duration of USVs, and altered the syllable composition in pups. Moreover, pups born to young fathers showed convergent vocal characteristics with a rich repertoire during postnatal development, while those born to aged fathers exhibited more divergent vocal patterns with limited repertoire. Principal component analysis in conjunction with clustering analysis demonstrated that pups from aged fathers deviated from typical trajectories of vocal development, which were considered as atypical individuals. Thus, our study indicates that advanced paternal age has a significant effect on offspring’s early vocal development. It is suggested that the trajectories of vocal development could be a useful marker of the NDD-like phenotype associated with the advanced paternal age. In addition, our comprehensive computational analysis described here is an effective approach to characterize the altered individual diversity relevant to neurodevelopmental disorders.

**One Sentence Summary:** Advanced paternal age affects vocal development in early postnatal mice, with more pups showing atypical developmental trajectories.

## Introduction

Human infant crying is an innate form of social communication^1,2^ that is used to attract attention from caregivers^3^ and affects cognitive control in adults^4^. It has been suggested that infant crying can serve as a marker of the behavioral and cognitive development and the altered crying features may indicate a risk for autism spectrum disorder (ASD)^5-7^ and other neurodevelopmental disorders (NDDs)^8,9^. Although the etiology of NDDs remains unclear, genetic, epigenetic, and environmental factors are likely to play an important role^10-14^. Recent epidemiological studies have shown a significant association between an advanced paternal age and the risk of NDDs in offspring^15-21^. Thus, we have investigated whether the advanced paternal age affects infant crying in mice and whether this phenotype may help to understand NDDs by providing predictive validity in mouse genetic studies.

Ultrasonic vocalizations (USVs) in rodents, particularly in mice, have mostly been investigated in relation to the neurobiology of vocal communication^22,23^. The separation of a pup from its mother and littermates induces the emission of USVs consisting of various sound elements (i.e., syllables) and the USVs can trigger maternal approach and retrieval behavior^24,25^. Neonatal USVs in mice could model some aspects of human infant cries^26^, and interestingly, exhibit gradual postnatal developmental changes in their acoustic features and compositions^27^. Mouse models for NDDs have been shown to exhibit various differences in USV parameters, including reduced number, higher or lower syllable frequency and shorter durations^28-31^. Previously, we reported that an advanced paternal age alters offspring’s behavioral phenotypes relevant to NDDs in mouse models^32,33^. It leads to deficiencies in the number, composition, and properties of USVs at postnatal day 6 (P6)^33^ in C57BL/6J mice, the most common inbred strain for behavioral and genetic analyses in preclinical studies.

Here, we comprehensively evaluated the developmental trajectory of early vocal communication in postnatal C57BL/6J mice derived from young and aged fathers, using semi-automatic, supervised, and unsupervised computational methods. Our data suggest that an advanced paternal age diversifies the developmental trajectories of vocalization, resulting in an increased proportion of atypical individuals.

## Results

Offspring were obtained by mating young (3-month-old) female C57BL/6J mice with either young (3-month-old) or aged (20-month-old) male C57BL/6J mice (YFO and AFO, respectively, **Fig.1A**). To record the USVs, each pup was separated one by one from its mother and littermates. The USV sonograms were transferred from the recorded acoustic waveforms and analyzed by three tools: a semi-automatic procedure (USVSEG)^34^, a supervised machine learning approach (VocalMat)^35^, and an unsupervised modeling algorithm (variational autoencoders, VAE)^36^ (**Fig. 1B**). The USVSEG requests manual syllable classification and noise inspection. To confirm and quantify the USV alterations without a bias from a human perspective, we also performed the VocalMat and VAE.

**Fig. 1.**
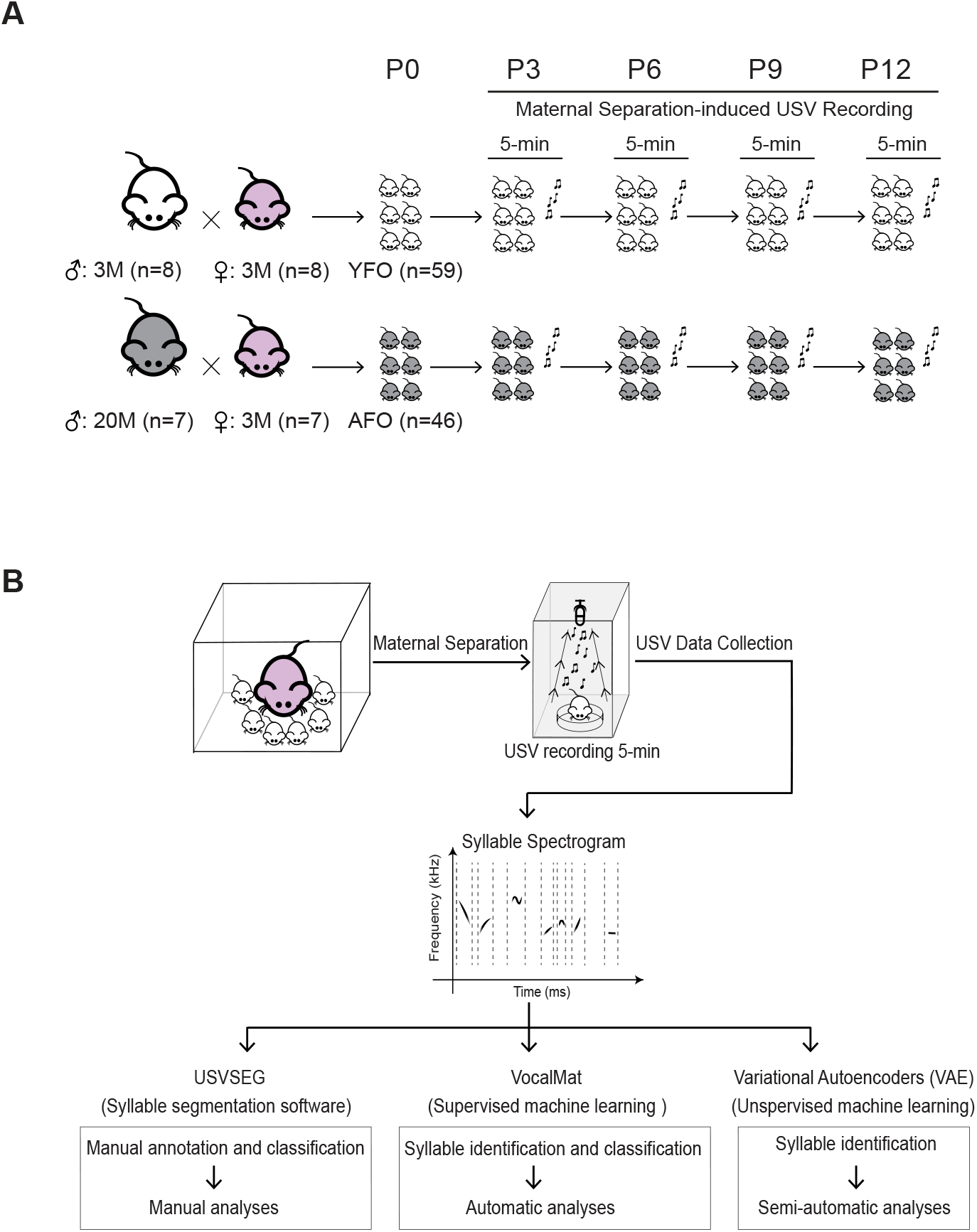
The experimental design and methods of syllable analyses (A) Workflow chart of the experimental design. Offspring were obtained by mating young C57BL/6J female mice with either young (3-month-old) or aged (20-month-old) C57BL/6J male mice (YFO and AFO groups, respectively). At P3, P6, P9 and P12, each pup was separated from its mother and littermates for a 5-min ultrasonic vocalization (USV) recording. **(B)** The maternal separation-induced USVs were processed and analyzed using USVSEG, VocalMat, and variational autoencoders (VAE).

### Advanced paternal age altered syllable properties and composition during development

The USVSEG revealed that the AFO group emitted a fewer USVs (fathers’ age, F (1, 412) = 55.902, *p* < 0.001; postnatal day, F (3, 421) = 7.339, *p* < 0.001; fathers’ age × postnatal day, F (3, 412) = 1.936, *p* = 0.123, mixed model) with shorter durations (fathers’ age, F (1, 412) = 52.173, *p* < 0.001; postnatal day, F (3, 421) = 22.695, *p* < 0.001; fathers’ age × postnatal day, F (3, 412) = 2.389, *p* = 0.068, mixed model) (**Fig. 2A-2B**). No significant difference was found in the maximum frequency or amplitude between the groups (**Fig. 2C-2D**). To further characterize call types, we classified syllables into 12 types (**Fig. 2E**) based on their spectrotemporal patterns^37^. Overall, the AFO group presented with a reduced number of syllables in three types (mixed model with post hoc comparison Tukey-Kramer test, see **Supplementary table 1** for results of mixed model). Also, this group of pups had shorter durations in one jump syllables and higher maximum amplitudes in two types of syllables (chevron and wave), but no significant difference was found in the maximum frequency (**Supplementary Fig. 1 and Supplementary Table 1**).

**Fig. 2.**
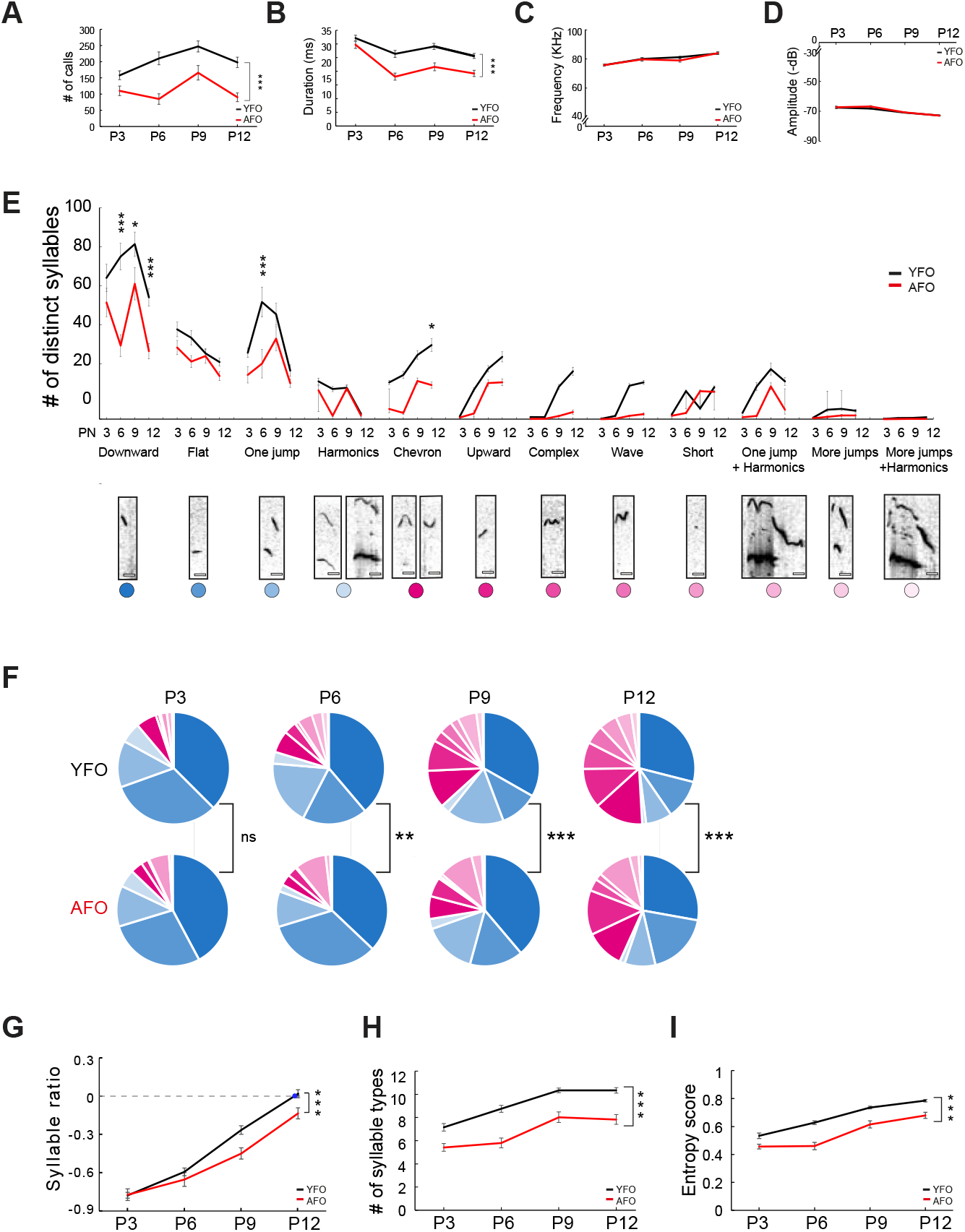
Advanced paternal age altered syllable properties and composition during development (data were obtained from USVSEG) (A) The number of calls in pups sired by young (3-month-old) or aged (20-month-old) male mice (YFO and AFO groups, respectively). *** *p* < 0.001 indicates a significant fixed effect of fathers’ age (mixed model). Data are shown as the mean ± standard error of the mean (SEM) for each group. (B) Syllable durations in the YFO and AFO groups. *** *p* < 0.001 indicates a significant fixed effect of fathers’ age (mixed model). Data are shown as the mean ± SEM for each group. (C) The syllable maximum frequencies in the YFO and AFO groups. Data are shown as the mean ± SEM for each group. (D) The syllable maximum amplitudes in the YFO and AFO groups. Data are shown as the mean ± SEM for each group. (E) Twelve distinct types of syllables^37^ were obtained. \**p* < 0.05, *** *p* < 0.001 indicate a significant difference between the two groups (Tukey-Kramer test, see Supplementary Table 1 for the results of mixed model). Data are shown as the mean ± SEM for each group. (F) Pie graphs illustrating the percentage composition of the 12 types of syllables from P3 to P12 in the YFO and AFO groups. Colors indicate the syllable types shown in (C). \*\**p* < 0.01 and *** *p* < 0.001 indicate a significant difference between the two groups (multivariate analysis of variance following Benjamini-Hochberg correction). (F) The syllable ratio was calculated as the difference between the number of pink spectrum syllables minus blue spectrum syllables and the number of pink spectrum syllables plus blue spectrum syllables, as follows: 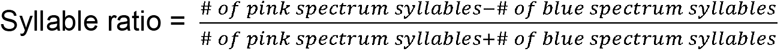 The blue dot indicates that the ratio reached zero. *** *p* < 0.001 indicates a significant fixed effect of fathers’ age (mixed model). Data are shown as the mean ± SEM for each group. (H) Developmental trajectories of syllable types in the YFO and AFO groups. *** *p* < 0.001 indicates a significant fixed effect of fathers’ age (mixed model). Data are shown as mean ± SEM for each group. (I) Entropy scores for syllables in the YFO and AFO groups. *** *p* < 0.001 indicates a significant fixed effect of fathers’ age (mixed model). Data are shown as the mean ± SEM for each group.

The syllable analysis showed that in the YFO group all types of syllables changed the percentage from P3 to P12: four types of syllables (downward, flat, one jump, and harmonics) decreased the percentage while other types of syllables increased the percentage. (**Supplementary Fig. 2**). We next analyzed the developmental transition of decreasing (blue spectrum colors, **Fig. 2F**) and increasing (pink spectrum colors, **Fig. 2F**) syllable types. The syllable types of blue spectrum colors were gradually replaced with those of pink ones from P3 to P12 in both YFO and AFO groups (**Fig. 2F**). While no significant difference was detected at P3, the syllable compositions significantly differed between the AFO and YFO groups from P6 to P12 (fathers’ age, P6: F (11, 93) = 2.687, *p* = 0.006; P9: F (11, 93) = 6.266, *p* < 0.001; P12: F (11, 93) = 4.150, *p* <0.001, multivariate analysis of variance [MANOVA]) (**Fig. 2F**). To investigate the dynamics of syllable composition during development, we used a syllable ratio to evaluate how quickly pups’ syllables transited from the initial period that the syllable proportion of blue spectrum color was enriched to the later period that the syllable proportions of blue and pink spectrum colors were equal (**Fig. 2G**). At P3, the syllable ratio was almost identical between the YFO and AFO groups. Although the YFO group showed the syllable ratio reaching to zero before P12, the AFO group did not (fathers’ age, F (1, 412) = 13.376, *p* < 0.001; postnatal day, F (3, 421) = 138.201, *p* < 0.001; fathers’ age × postnatal day, F (3, 412) = 2.612, *p* = 0.051, mixed model).

Finally, we addressed the syllable diversity. While the YFO group emitted USVs that contained an increasing number of syllable types with age, the AFO group produced USVs with a significantly smaller number of syllable types from P3 to P12 (fathers’ age, F (1, 412) = 100.251, *p* < 0.001; postnatal day, F (3, 421) = 35.638, *p* < 0.001; fathers’ age × postnatal day, F (3, 412) = 1.086, *p* = 0.355, mixed model) (**Fig. 2H**). To quantify the syllable properties, entropy score, which ranged from 0 to 1, was calculated to indicate production uniformity^27^ (**Fig. 2I**). The score approached 1 when a pup uniformly (or diversely) produced all the syllable types and approached 0 when a pup preferred to produce only one syllable type (limited repertoire). The entropy score increased continuously in both groups during development. However, the AFO group consistently showed lower entropy scores than the YFO group across all postnatal stages (fathers’ age, F (1, 412) = 86.025, *p* < 0.001; postnatal day, F (3, 421) = 77.095, *p* < 0.001; fathers’ age × postnatal day, F (3, 412) = 2.207, *p* = 0.087, mixed model).

The above data demonstrate that advanced paternal age reduced the number and duration of specific call types, delayed development of syllable compositions, and limited syllable repertoire in offspring. Thus, the AFO group had overall a limited vocalization repertoire.

### Advanced paternal age increased postnatal USV variation in offspring

To compare the developmental trajectories among individual mice, we applied principal component analysis (PCA) to extract the syllable features of the USVSEG data (**Fig. 3A**) using five syllable parameters (**Fig. 3B**). At P3, the circles comprising 90% of the offspring in both groups were wide and almost overlapped, indicating that individual P3 pups were diverse in vocal repertoire regardless of paternal age. From P6 to P12, the YFO group showed a developmentally convergent pattern with gradually smaller circles including 90% of YFO (gray circles in **Fig. 3A**). In contrast, a relatively wider individual difference remained in the AFO group, even at P12 (pink circles in **Fig. 3A**). This longitudinal analysis clearly indicated the different developmental trajectories of the YFO and AFO groups (fathers’ age, F (1, 418) = 106.864, *p* < 0.001, postnatal day, F (3, 421) = 20.132, *p* < 0.001; fathers’ age × postnatal day, F (3, 412) = 2.412, *p* = 0.066, MANOVA).

**Fig. 3.**
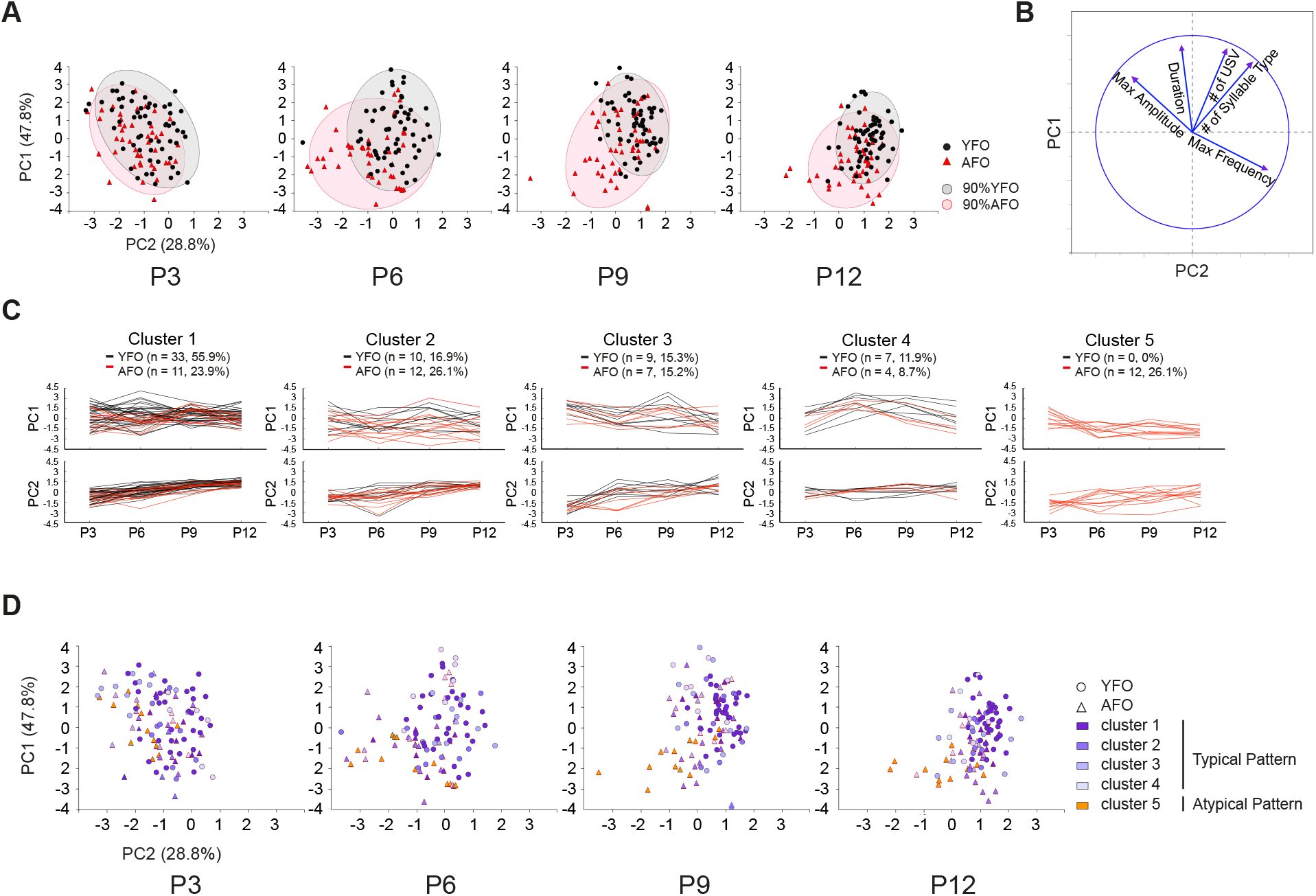
Advanced paternal age influenced individual trajectories of vocal development and resulted in an increased number of atypical individuals (data were obtained from USVSEG) (A) Principal component analysis (PCA) summarized the patterns of individual pups sired by young (3-month-old) or aged (20-month-old) male mice (YFO and AFO groups, respectively) from P3 to P12. Each point represents a syllable pattern of a single pup. Shorter distances between points indicate greater similarity in the syllable patterns. The grey and pink circles contain 90% of the population of the YFO and AFO groups, respectively. (B) The loading plot of the PCA shows correlations between the original ultrasonic vocalization (USV) parameters and two principal components. (C) Clustering analysis separated individual offspring into five clusters according to PC1 and PC2. (D) Based on the cluster analysis according to PC1 and PC2, the pups of Clusters 1–4 were classified as typical individuals and the pups of Clusters 5 were classified as atypical individuals. Typical and atypical pups are marked on the PCA spatial map.

To better understand the individuality of USVs during development, clustering analyses were performed using Gaussian mixed models (GMMs) along with the Akaike information criterion (AIC) (**Supplementary Table 2**). The number of clusters was objectively determined using the minimum AIC. PC1 and PC2 were loaded together to apply clustering analyses and were clustered into five different clusters that shared a common pattern (**Fig. 3C**). The different cluster patterns were apparent between the YFO and AFO groups; the YFO group was separated into four clusters (see black lines) and enriched in Cluster 1 (55.9%), while the AFO group was separated into five clusters (see red lines) and was enriched in Cluster 5 (26.1%, χ2 = 22.993, *p* < 0.001, chi-squared independence test). A similar phenotype was observed in the additional cluster analyses using other USV parameters. The number of calls was positively correlated with syllable duration (YFO: Pearson’s correlation coefficient *r* = 0.512, *n* = 59, *p* < 0.001; AFO: *r* = 0.547, *n* = 46, *p* < 0.001) and these two factors were clustered and separated into five clusters (**Supplementary Fig. 3A**). The developmental trajectories of the number of calls and duration observed in the YFO group were distributed among four clusters and enriched in Cluster 1, while those in the AFO group were distributed among five clusters and enriched in Cluster 3 to 5 (**Fig. 3C**). The cluster patterns (i.e., the proportion of individuals in each cluster) significantly differed between the YFO and AFO groups (χ2 = 14.810, *p* = 0.005, chi-squared independence test).

Next, we clustered the developmental trajectories of the number of syllable types into five clusters (**Supplementary Fig. 3B**), in which the YFO group was again enriched in Cluster 1, while the AFO group was in Clusters 3 to 5. There was a significantly difference in the cluster patterns between the YFO and AFO groups (χ2 = 29.642, *p* < 0.001, chi-squared independence test). Finally, to examine the individual developmental patterns of syllable properties, we clustered the entropy scores. Again, the cluster distribution significantly differed between the YFO and AFO groups (χ2 = 31.304, *p* < 0.001, chi-squared independence test). The YFO group occupied three clusters and was dominant in Cluster 1, whereas the AFO group occupied five clusters and was dominant in Cluster 3 to 5 (**Supplementary Fig. 3C)**. Therefore, the longitudinal developmental patterns significantly differed between the YFO and AFO groups, with some patterns unique only to individuals in the AFO group.

A unique trajectory pattern (i.e., Cluster 5) was observed in the clustering analysis (Fig. 3C). Because this cluster only comprised with AFO, we evaluated the pups in Cluster 5 as “atypical” individual ones. Conversely, pups in Clusters 1 to 4 were judged to be “typical” ones. These typical and atypical pups were labeled with purple and yellow colors in the PCA data (**Fig. 3D and Supplementary Fig. 4**). From P3 to P12, the distance among the typical pups gradually decreased. Although the pups were mixed at P3, the typical and atypical pups gradually diverged as they developed (**Fig. 3D)**. The convergent phenotype was more apparent in the YFO group because this group contained more typical individuals (**Supplementary Fig. 4**). These data demonstrate that atypical individuals with a unique pattern were originated from 26.1% of the AFO group.

**Fig. 4.**
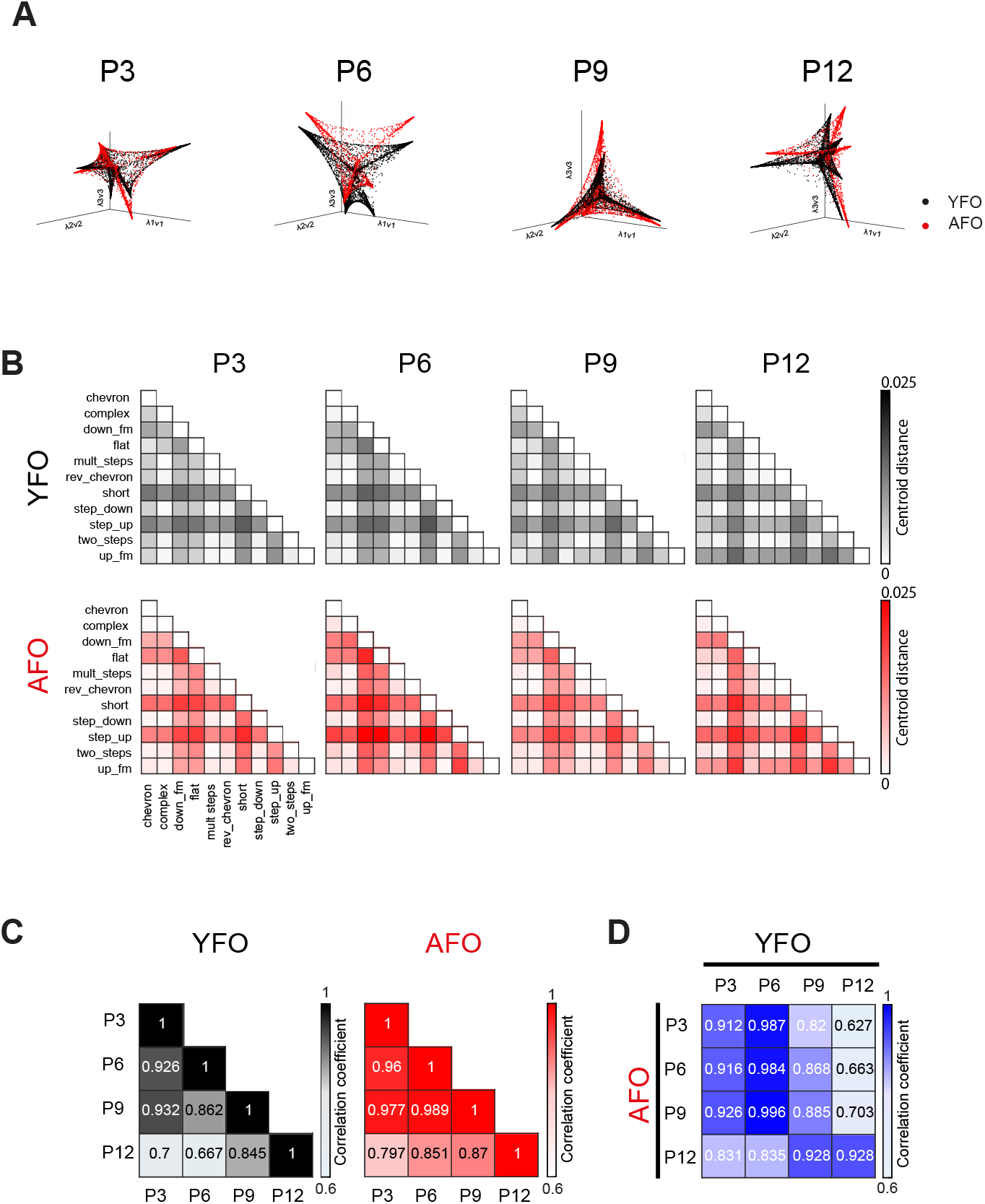
Advanced paternal age led to a different three-dimensional distribution of syllable composition (data were obtained from VocalMat) (A) Syllables from pups sired by young (3-month-old) or aged (20-month-old) male mice (YFO and AFO groups, respectively) were mapped in a diffusion Map after classification from P3 to P12. (B) The pairwise distance between the centroids of syllable types within each manifold structure. (C) The matrix of Pearson’s correlation within groups. The number indicates the correlation coefficient. (D) The matrix of Pearson’s correlation between groups. The number indicates the correlation coefficient.

### Advanced paternal age altered the postnatal development of syllable composition in pups

To confirm and quantify the alteration of syllable composition without a bias from a human perspective, we next applied a supervised machine learning method, VocalMat, for the automated detection and classification of syllables^35^. After classifying the syllables into 11 categories, VocalMat visualized the three-dimensional probability distribution of the syllables using diffusion maps, which showed a dynamic change from P3 to P12 (**Supplementary Fig. 5**). Differences in syllable composition between the YFO and AFO groups were also visually verified (**Fig. 4A)**. To quantitatively evaluate measure these differences, the pairwise distance between the centroids of syllable types within each manifold structure was computed as a heatmap (**Fig. 4B**). For a summary and direct comparison of the manifolds between the groups across the four postnatal days, matrix correlations were calculated within and between groups (**Fig. 4C, D**). The three-dimensional syllable compositions underwent gradual changes within both groups and the highest similarity was observed between P6 YFO and P9 AFO when comparing the two groups. These results confirm that the AFO group emitted USVs with different syllable compositions and developmental delay.

**Fig. 5.**
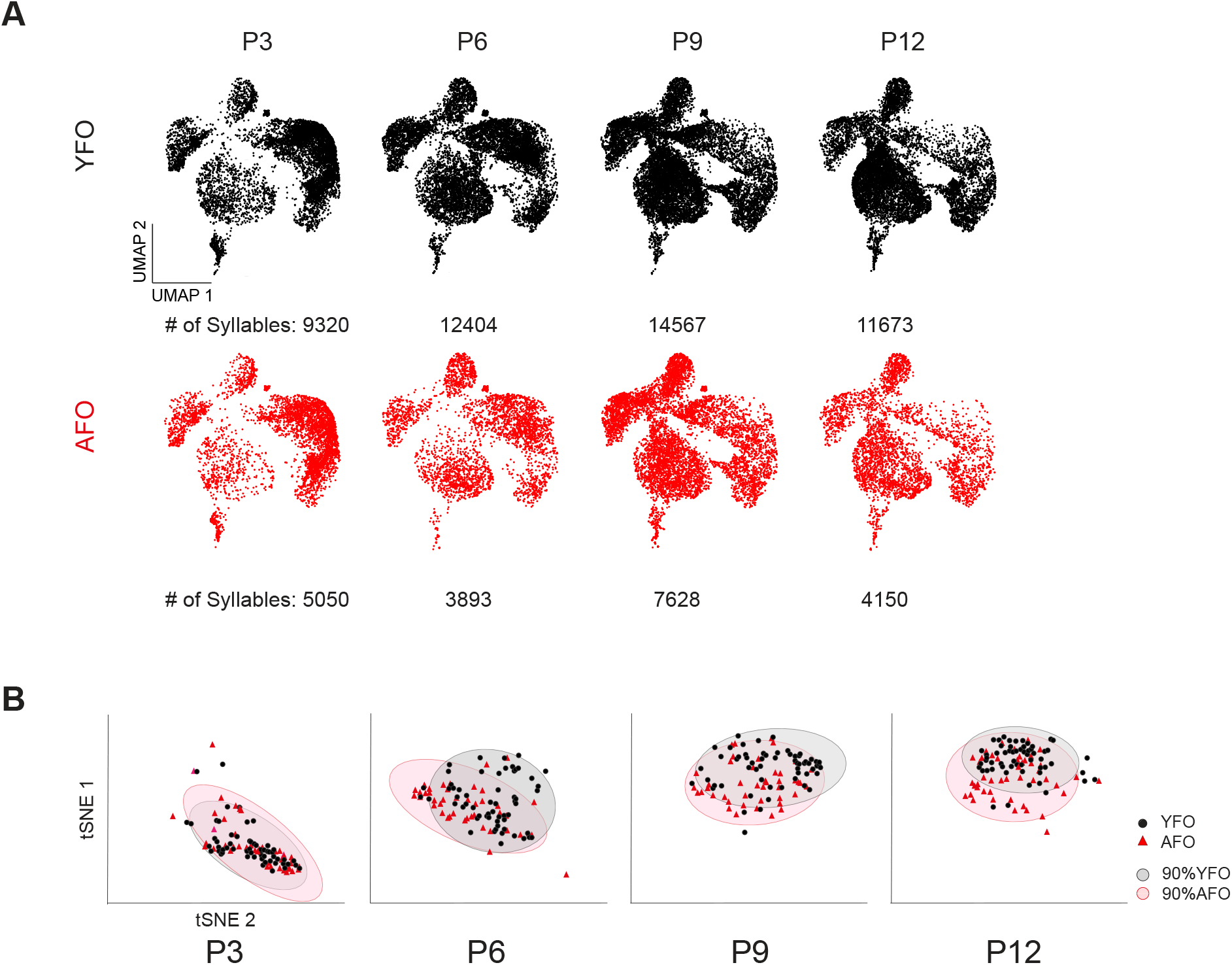
Variational autoencoders confirmed that the advanced paternal age affected the convergent pattern of syllable development (data were obtained from variational autoencoders) (A) Variational autoencoders (VAE) mapped syllables from pups sired by young (3-month-old) or aged (20-month-old) male mice (YFO and AFO groups, respectively). Each point in this latent space represents a single syllable. The distance between points indicates neighbor similarity, with closer points being more similar. (B) Based on VAE, t-SNE summarized the patterns of individual pups from the YFO and AFO groups from P3 to P12. Each point represents the syllable pattern of one pup. Shorter distances between points indicate greater similarity in syllable patterns. Grey and pink clusters contain 90% of the population of the YFO and AFO groups, respectively.

### VAE may support the convergent pattern of syllable development

To objectively identify important USV variations, we further applied an unsupervised modeling algorithm, VAE^36^, to characterize subtle changes without a human bias of syllable classification. Each point representing one syllable was mapped to the inferred latent space and visualized (**Fig. 5A**). A unique syllable distribution at each developmental stage in both the YFO and AFO groups seems to show that vocalization patterns developed through a dynamic process, which is consistent with our results from USVSEG and VocalMat. However, it is difficult to quantify and compare these syllable patterns directly between the YFO and AFO groups. In the AFO map, some areas with lower densities were noted. Therefore, we applied t-distributed stochastic neighbor embedding (t-SNE) to the VAE data to reduce the dimensions and summarize the patterns to allow the comparison of individual differences (**Fig. 5B**). Again, different patterns were identified (fathers’ age, F (1, 404) = 35.803, *p* < 0.001, postnatal day, F (3, 404) = 196.583, *p* < 0.001; fathers’ age × postnatal day, F (3, 404) = 3.872, *p* = 0.009, MANOVA). The post-hoc test revealed that the differences appeared from P6 to P12 (P3: F (1, 404) = 0.020, *p* = 0.888; P6: F (1, 404) = 10.423, *p* = 0.001; P9: F (1, 404) = 17.310, *p* < 0.001; P12: F (1, 404) = 19.773, *p* < 0.001, F test). The vocal repertoires of the YFO group demonstrated developmental convergence despite different trajectories, whereas those of the AFO group were relatively wide scattered during development.

The consistent results of the USVSEG and VAE findings were as follows: 1) the inferred spatial positions of the YFO and AFO groups were clearly similar at P3 but became significantly different in later developmental stages; and 2) a relatively divergent pattern of syllable development was observed in the AFO group, in contrast to the convergent pattern noted in the YFO group.

### Advanced paternal age affected the body weight gain

To explore the implications of vocal communication in the physical development of neonatal mice, we measured the body weight after each USV recording. Lower body weight gain was consistently observed in the AFO group than in the YFO group (fathers’ age, F (1, 412) = 35.888, *p* < 0.001; postnatal day, F (3, 421) = 1842.759, *p* < 0.001; fathers’ age × postnatal day, F (3, 412) = 2.196, *p* = 0.088, mixed model) (**Fig. 6A**). To determine whether syllable development was associated with body weight gain, Pearson correlation was applied and visualized as a heatmap (**Fig. 6B**). Unexpectedly, a greater number of syllable parameters was significantly correlated with body weight in the AFO group than in the YFO group. Further, in the YFO group, most syllable parameters showed significant correlations with body weight on P6; but in the AFO group, P9 was the equivalent developmental stage. Thus, the AFO group displayed lower body weight gain and a delayed postnatal time point, although a specific correlation between body weight and USVs was difficult to evaluate.

**Fig. 6.**
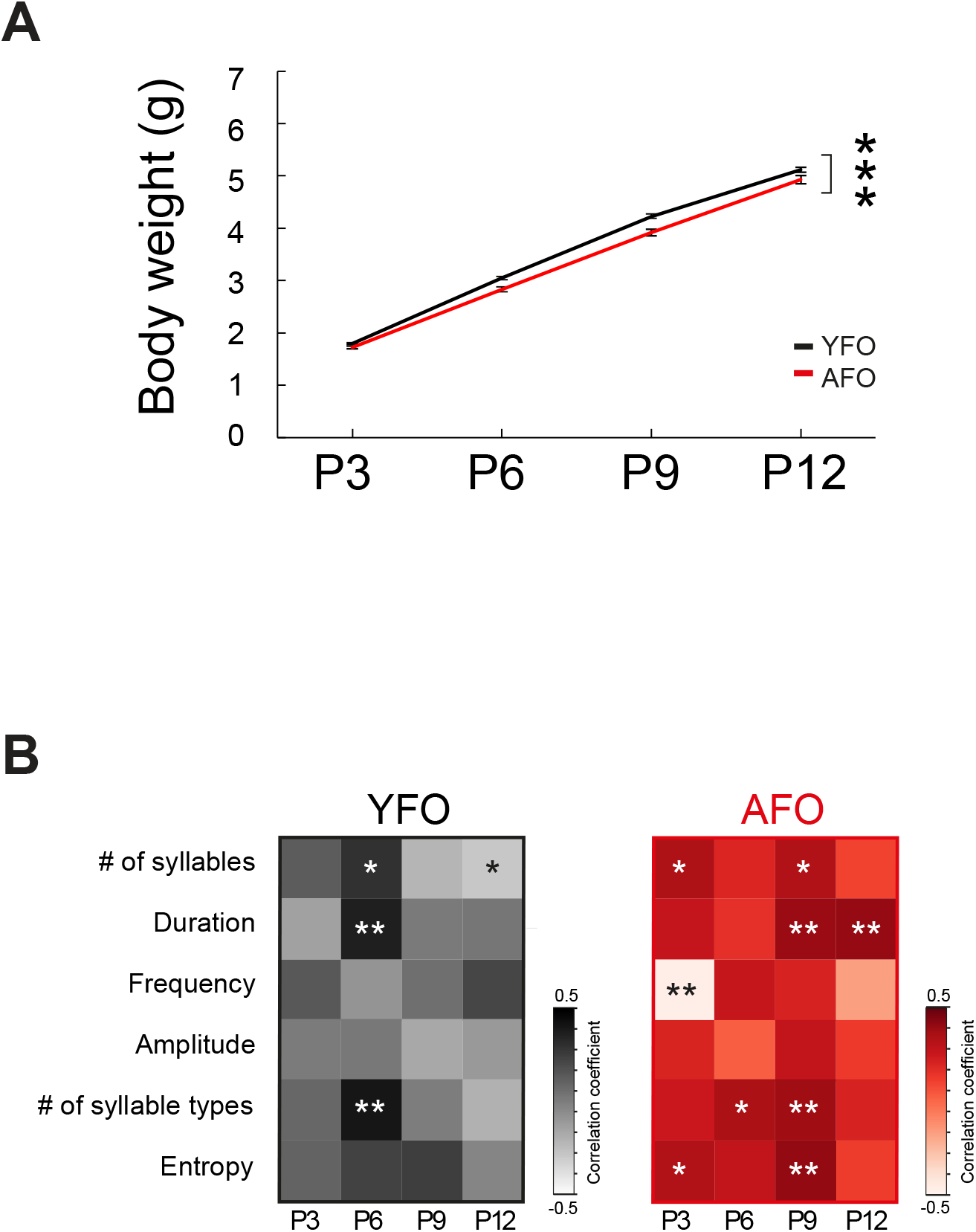
Advanced paternal age affected the body weight gain (A) Body weight of pups sired by young (3-month-old) or aged (20-month-old) male mice (YFO and AFO groups, respectively). *** *p* < 0.001 indicates a significant fixed effect of fathers’ age (mixed model). Data are shown as the mean ± standard error of the mean (SEM) for each group. (B) Pearson’s correlation between body weight and ultrasonic vocalization (USV) parameters that were obtained from USVSEG was calculated and presented from P3 to P12 as a heatmap. \**p* < 0.05 and \*\**p* < 0.01 indicate a significant correlation between the two variables.

## Discussion

Using the three sophisticated computational analyses, we report that an advanced paternal age diversifies the developmental trajectories of USVs in mice at early postnatal stages. One of our main findings was that the advanced paternal age causes alterations in early vocal behavior and increases the number of offspring with atypical developmental patterns. This phenotype recapitulates clinical evidence that an advanced paternal age is a risk factor for the atypical development observed in children with NDDs^15-17^, and suggests that the effect of advanced paternal age could be detected in early infancy.

Previous studies reported that USVs from mouse pups may be analogous to crying in human infants and are thus one of the few methods used to investigate behavioral development during the early postnatal period ^38,39^. In our study, the YFO group emitted diverse types of syllables during the postnatal period, and this finding is consistent with that of a previous study demonstrating normal syllable development in CBA/CaJ mouse pups^27^. Compared to the YFO group, the AFO group emitted a narrower spectrum of syllable types with a reduced number of syllables, which is reminiscent of the poorer vocal repertoire in some ASD patients^40^. A reduction in syllable repertoire has also been described in genetic NDD models, such as *Cd157* KO mice^41^ and *Tbx1* heterozygous mice^42,43^. Additionally, we observed an altered composition of syllables in the AFO group, which has previously been reported in other models, such as the *Reelin* mutant^44^, *fmr1* knockout^45^, *ScSn-Dmd*^*mdx*^*/J* mutant^46^ and *Tbx1* heterozygous mice^42,43^. Thus, our model of advanced paternal age showing impairments in syllable properties and composition also fits for the NDD-like phenotype.

There is a consensus that early childhood development typically follows a series of developmental milestones within a similar age range. However, 10%-17.8% of children demonstrate mild to severe developmental delays, including NDDs, where delayed and/or atypical development has been considered an early warning sign^47-50^. Our mouse model revealed that individual pups in the AFO group showed deviated patterns becoming most significantly different at P12, while the YFO group exhibited typical development of vocal repertoires gradually converging with age. Additionally, 26.1% of the AFO group displayed a unique developmental pattern and was identified as atypical individuals. This phenotype suggests that the impact of advanced paternal age is detectable in offspring during an early neonatal period in mice. We notice that the unique pattern of AFO was not only displayed in the clustering analysis based on PCA data (**Fig. 3C**), but also was showed in other cluster analyses based on syllable parameters (**Supplementary Fig. 3**). These results may elaborate the heterogeneity of NDDs and mirror children with NDDs showing a variety of atypical developmental behaviors^51-53^, including non-uniformly impairment that becomes more distinct with age^54-56^. It would be interesting to elucidate the underlying mechanisms and determine whether those atypical pups show behavioral deficiencies in adulthood stage in future studies. A limitation here is that no standard criteria can be used to determine the typical and atypical individuals in neonatal USVs. Therefore, we do not rule out the possibility that atypical individuals also exist in the YFO group.

Since sensitivity to behavioral signs is low during early development and the phenotypic characteristics of NDDs are complex and heterogeneous in nature, early diagnosis of NDDs is challenging at the clinical level. Recently, dimensional and/or clustering approaches have been applied to identify developmental changes and to screen individuals in general or at-risk populations^57^. Moreover, a mouse model of 16p11.2 copy number variation, a risk for NDDs, has shown that dimensional features of peripubertal social behaviors can be computationally predicted by developmental trajectories of neonatal USVs^58^. In the present study, PCA and clustering analysis were integrated to extract the complicated features of USVs and identify individual pups with atypical USV patterns. These approaches may provide a translational perspective to capture the underlying heterogeneity of NDDs in the population at large and offer valuable suggestions for early diagnosis and intervention in the children with NDDs.

Epidemiological studies have consistently demonstrated that the advanced paternal age causes smaller fetuses in gestational age, as well as lower body weight^59,60^. In humans, the association between physical growth, including body weight gain, and neurodevelopmental delay has been examined in general populations of term-born infants; it was found that infants with poor growth in early infancy are at an increased risk of neurodevelopmental delay^61,62^. We noticed that during the first two postnatal weeks, the AFO group exhibited lower body weight gain, which was also observed in a rat model of advanced paternal age^20^. We also observed a similar tendency in our previous study, in which we only recorded USV and body weight at P6^33^. There are two possible explanations for the lower gain of body weight in the AFO group. First, the advanced age of father could directly cause developmental delay of the physical conditions in offspring. However, we observed a significantly developmental delay in USV parameters (Fig. 2–3) but robust deviation in individual difference or syllable repertoire complexity (Fig. 4-5) in the AFO group. The group differences of USVs, therefore, could not simply reflect the physical developmental delays. The genetic and/or epigenetic factors associated with the advanced paternal age might trigger lower body weight gain affecting on the other factors irreversibly during postnatal development.

Another possibility is that body weight gain is influenced by altered maternal behaviors due to differences in the communication of the offspring including USVs. It is well known that pup USVs have communicative significance^63-67^, which is crucial for maternal care behaviors, preserves the social bonds between mother and infant, and is essential for the healthy development of offspring^68,69^. Previous studies have shown that USVs of *Tbx1* heterozygous pups do not induce maternal approach behaviors^42,70^. More work is needed to further determine whether USVs emitted by AFO elicit less maternal approach behavior and thus lower body weight. Although underlying mechanisms are not fully understood, an advanced paternal age affects the physical condition and neurodevelopment of pups, possibly via a combination of genetic, epigenetic, and environmental factors.

Although approaches to analyze USV have advanced dramatically during the past decade, it is undeniable that any method has its limitations. The semi-automatic approach (USVSEG) provides a reliable segmentation performance for USV syllables^34^, whereas the manual syllable classification and noise inspection were time-consuming and affected by human bias. VocalMat, a supervised machine learning method, is able to automatically detect and classify USVs^35^ but still has a room improved for its accuracy. On the other hand, an unsupervised machine learning algorithm (VAE) can capture data variability but must be trained on each dataset, with the limitation that mouse USV syllables cannot be categorized into distinct types^36^.

In the current study, we applied a composite of these three approaches, understanding the strengths and limitations of each, and collected reliable data from different angles. USVSEG demonstrated the detailed alterations of acoustical features and composition that were obtained after a half year’s work (Fig. 2-3). VocalMat contributed to the high-throughput detecting difference and similarity of syllable repertoire within days (Fig. 4 and Supplementary Fig. 5) although it may not cover all the syllable categories. VAE explored the developmental transition of syllable variability and individual difference with minimizing human bias although the bulk of syllable difference was relatively ambiguous between groups (Fig. 5). Even though these different approaches do not share the same computer algorithms, we were able to reach a unanimous conclusion from these comprehensive USV analyses; USVs in the AFO group showed altered composition and different developmental trajectories. Technology advances in extracting information from USVs can solve shortcomings of existing methods and, most importantly, unify the criteria to identify translational relevant traits.

Our findings provide evidence for the hypothesis that the advanced paternal age expands distinct individuals reflecting “neurodiversity” in offspring. Moreover, as in modern societies the age when individuals give birth is increasing, advanced paternal age may represent a risk factor for neurodevelopmental disorders^71^. In these cases, epigenetic changes are the most likely mechanisms to underly this phenomenon^72^. Another possibility is that advanced paternal age causes the accumulation of spontaneous *de novo* mutations^73,74^, possibly leading to neurodevelopmental disorders. Last but not least, increased longevity occurred during human evolution. The investigation of sophisticated developmental traits through epigenetic inheritance is necessary to account for the process of civilization.

## Acknowledgements

We are grateful for the financial support of this research by KAKENHI in Innovative Areas (Grant Number 16H06530) from the Ministry of Education, Culture, Sports, Science, Technology and from the Japan Agency for Medical Research and Development (AMED, Grant Number JP21wm0425003) to N.O. and funds to N.H. (R01DC015776, NIDCD/NIH; R21HD105287, NICHD/NIH: R01MH099660, NIMH/NIH).

## Author contributions

Study design: L. M. and N. O.

USV collection and analysis: L. M.

Contributed analytic tools: K. K., R. O. T

USV data analysis: L. M., R. K., H. I., T. M., R. O. T., F. K.

Technique for USV recording: L. M. and R.K.

Draft the paper: L. M., H. I., V.T., N. H., N. O.

Discussed and interpreted the results: all authors

## Declaration of interests

The authors declare no competing interests.

## Data and code availability

All data in this paper will be shared by the lead contact upon request. The original code and any additional information required to reanalyze the data reported in this paper are available from the lead contact upon request.

**Supplementary Table 1:** The results of mixed model

**Supplementary Table 2:** The AIC number according to the number of clusters

**Supplementary Fig. 1**. Advanced paternal age affected the acoustic features of distinct types of syllables (data were obtained from USVSEG)

(A) The duration of 12 distinct types of syllables in pups sired by young (3-month-old) or aged (20-month-old) male mice (YFO and AFO groups, respectively). \*\**p* < 0.01 indicate a significant difference between the two groups (Tukey-Kramer test, see Supplementary Table 1 for the results of mixed model). Data are shown as the mean ± standard error of the mean (SEM) for each group.

(B) The maximum frequencies of 12 distinct types of syllables in the YFO and AFO groups. see Supplementary Table 1 for the results of mixed model. Data are shown as the mean ± SEM for each group.

(C) The maximum amplitudes of 12 distinct types of syllables were in the YFO and AFO groups. \*\*\**p* < 0.001 indicate a significant difference between the two groups (Tukey-Kramer test, see Supplementary Table 1 for the results of mixed model). Data are shown as the mean ± SEM for each group.

**Supplementary Fig. 2**. In the YFO group, 12 types of syllables changed the percentages from P3 to P12 and were classified into two categories: decreasing (blue spectrum colors) and increasing (pink spectrum colors) percentages.

(A) –(D) In the YFO group the percentage of four types of syllables (downward, flat, one jump, and harmonics) decreased from P3 to P12 (downward: postnatal day, F (3, 235) = 9.536, *p* < 0.001, one-way repeated analysis of variance [ANOVA]; flat: postnatal day, F (3, 235) = 39.665, *p* < 0.001, one-way repeated ANOVA; one jump: postnatal day, F (3, 235) = 12.069, *p* < 0.001, one-way repeated ANOVA; harmonics: postnatal day, F (3, 235) = 10.669, *p* < 0.001, one-way repeated ANOVA)

(E)–(L) In the YFO group the percentage of eight types of syllables (chevron, upward, complex, wave, short, one jump + harmonics, more jumps, and more jumps + harmonics) increased from P3 to P12 (chevron: postnatal day, F (3, 235) = 29.039, *p* < 0.001, one-way repeated ANOVA; upward: postnatal day, F (3, 235) = 49.060, *p* < 0.001, one-way repeated ANOVA; complex: postnatal day, F (3, 235) = 69.168, *p* < 0.001, one-way repeated ANOVA; wave: postnatal day, F (3, 235) = 57.123, *p* < 0.001, one-way repeated ANOVA; short: postnatal day, F (3, 235) = 14.819, *p* < 0.001, one-way repeated ANOVA; one jump + harmonics: postnatal day, F (3, 235) = 9.869, *p* < 0.001, one-way repeated ANOVA; more jumps: postnatal day, F (3, 235) = 8.530, *p* < 0.001, one-way repeated ANOVA; more jumps + harmonics: postnatal day, F (3, 235) = 3.342, *p* = 0.02, one-way repeated ANOVA).

**Supplementary Fig. 3**. Advanced paternal age affected individual development longitudinally (data were obtained from USVSEG)

(A) Clustering analysis separated the individual offspring into five clusters according to the number of calls and syllable duration.

(B) Clustering analysis separated the individual offspring into five clusters according to the number of syllable types.

(C) Clustering analyses separated the individual offspring into five clusters according to the entropy.

**Supplementary Fig. 4**. Summary of individual traces at P3 and P12.

Based on Fig 3. (D). Each point represents the syllable pattern of one pup. P3 and P12 pups are indicated by transparent and opaque symbols, respectively. The trajectories from P3 to P12 for each pup are indicated by gray lines.

**Supplementary Fig. 5**. VocalMat confirmed that the advanced paternal age led to a different syllable composition

(A) VocalMat classified the syllables into 11 types and mapped them in a diffusion map from P3 to P12.

(B) According to (A), syllables were identified in the YFO and AFO groups.

## STAR Methods

### Animals

All experimental procedures were approved by the Ethics Committee for Animal Experiments of Tohoku University Graduate School of Medicine (#2014-112) and the animals were treated according to the National Institutes of Health Guidance for the Care and Use of Laboratory Animals. Briefly, 20-months-old male C57BL/6J mice were used as aged fathers because 18–24 months of age in mice correlates to 56–69 years of age in humans^75^, which meets the definition of “aged”. Eight young (3-month-old) and seven aged male mice were crossed with young (3-month-old) virgin female C57BL/6J mice. After mating, each female mouse was separated from the male mouse and isolated to minimize possible confounding factors related to offspring behavior. In this study, 59 offspring from young fathers and 46 offspring from aged fathers were used. Our exploratory preliminary analysis did not determine statistically significant effect for the sex. This result is consistent with previous studies^31,76^. Therefore, the sex effect was excluded from our analysis and both male and female pups were used. Offspring that died during the experimental period were excluded from this study (mortality rate, YFO group: 4.8% [3/62] vs. AFO group: 8% [4/50]). At P3, each offspring was tattooed using the Aramis Animal Microtattoo System (Natsume Co., Ltd., Tokyo, Japan) for individual recognition after the USV test (described below). The average litter size and number of pups did not significantly differ between the YFO (7.75 ± 1.16) and AFO (7.14 ± 1.86) groups (*t* = 0.744, *p* = 0.474, *t*-test). All animals were housed in standard cages in a temperature-and humidity-controlled room with a 12-hour light/dark cycle (lights on at 08:00) and had free access to standard laboratory chow and tap water.

### USV recordings

According to previously described protocols^30,32,33,77^, each pup was separated from its mother and littermates, placed on a transparent plastic dish with woodchip bedding, and transferred to a sound-attenuating chamber for USV recordings at P3, P6, P9 and P12. An ultrasound microphone (Avisoft-Bioacoustics CM16/CMPA) was inserted through a hole in the middle of the cover of the chamber, approximately 10 cm above the offspring, to record vocalizations. The recorded vocalizations were transferred to an UltraSound Gate 416H detector set (Avisoft Bioacoustics, Germany) at 20–125 kHz. After a 5-min recording session, the body weight of the pups was measured and they were returned to their home cage. This procedure was repeated until all pups had been recorded. The room temperature was maintained at 22°C.

### Semi-automatic analysis of syllable properties

Acoustic waveforms were processed using USVSEG, a GUI-based MATLAB (MathWorks Inc., MA, USA) script, originally developed for segmenting USVs emitted by rodents^34,37^. Briefly, the script computed the spectrograms from each waveform (60 s/block), applied a threshold to remove noise from the signal, and detected syllables within a frequency range of 20–120 kHz. The segmentation criteria for identifying syllables were as follows: a minimum gap of 10 ms should separate two syllables and each syllable should have a minimum duration of 2 ms. This script segmented each syllable and exported them as individual jpeg files. The duration, peak frequency (frequency at maximum amplitude), and maximum amplitude of each syllable were calculated automatically. Based on previous published criteria^37^, by visual inspecting jpeg files, segmented syllables were manually classified into 12 types or excluded as noises (false positive). The pink/blue syllable ratio was calculated as the difference between the number of pink spectrum syllables minus blue spectrum syllables and the number of pink spectrum syllables plus blue spectrum syllables, as follows:

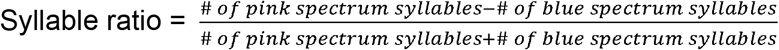

### Supervised automatic syllable classification

The supervised learning software VocalMat was used for the automatic detection and classification of syllables. Following the fixed threshold and parameters of the original published method for syllable segmentation, syllables were classified into 12 categories (11 syllable types + noise)^35^. VocalMat then visualized the syllable distribution using diffusion maps to reduce the 11 dimensions to three dimensions. The pairwise distance between the centroids of syllable types within each group at each stage was calculated and output as matrices using MATLAB.

### Unsupervised modeling and characterization of vocalizations

The VAE, which is an unsupervised learning method for extracting essential features without any instructional input, was used to characterize the recorded vocalizations. We used the VAE to reduce the dimensions of the raw data and quantify subtle changes in behavioral variability to preserve as much information as possible, according to an established procedure^36^. The VAE algorithm, which was implemented in PyTorch (v1.1.0), was trained to maximize the evidence lower bound as the standard setting. Because the audio segmentation method implemented in the original program detected large amounts of noise and was unable to detect some subtle syllables, we decided to use USVSEG for audio segmentation prior to using VAE. The syllables were detected and segmented by USVSEG using the same parameters as those mentioned above. After training, a latent map of all syllables was created using a uniform manifold approximation projection. To visualize the differences between individuals, t-SNE was applied to reduce the dimensions for each pup based on the maximum mean discrepancy (MMD)^36^. We excluded pups that emitted two or fewer calls (two in the AFO group at P3, three in the AFO group at P6, and three in the AFO group at P9) from the VAE analysis because the MMD cannot be applied in such datasets.

### Statistical analysis

A mixed model with the fixed effects (father’s age and postnatal day), random effects (litter), and repeated measures was applied to examine the statistical significance of overall syllable parameters, syllable ratio, number of syllable types, entropy, and body weight. To detect the statistical significance of number, duration, frequency, and amplitude of 12 distinct sylalbles, the mixed model with the fixed effects (father’s age, postnatal day, and syllable type) was performed. Tukey-Kramer test post hoc test was applied if interaction was significant. MANOVA was performed to detect differences of syllable composition in each postnatal day with effect of fathers’ age. The differences of principal components and t-SNEs between the YFO and AFO groups were detected using MANOVA with the father’s age and postnatal day as two independent effects. The post-hoc comparison was performed using F test if the interaction was significant. The Pearson’s correlation coefficient was used to determine the correlation between body weight and USV parameters.

The entropy score was calculated using the information theory toolbox in MATLAB. We excluded some offspring (one from the YFO group and one from the AFO group at P3, three from the AFO group at P6, and two from the AFO group at P9) whose total number of syllables and number of syllable types produced in 5 minutes was insufficient (three or fewer syllables and only one syllable type) from the entropy calculation. To understand the development at the individual level, clustering analysis with GMMs was applied, where the data dimension of eight corresponded to the number and duration of syllables at the four time points. We fitted GMMs with diagonal covariance Gaussian components using the MATLAB function *fitgmdist*. The number of clusters was determined by minimizing the AIC^78,79^. Based on the fitted GMM, each individual mouse pup was categorized into a cluster with the maximum posterior probability. The chi-square independence test was subsequently applied to determine whether the cluster distribution was significantly different between the two groups. A PCA was performed to objectively characterize the typical syllable patterns of individual offspring. In the present study, syllable data, including the syllable number, number of syllable types, duration, maximum frequency, and maximum amplitude, were inputted to the PCA to generate principal components. For all comparison, the significance level was set at α = 0.05. JMP13 Pro software (SAS Institute, Cary, NC, USA) was used for statistical analyses. Data are presented as mean ± standard error of the mean for each group.

## Notes

### Competing Interest Statement

The authors have declared no competing interest.

## References

1. Acebo, C., and Thoman, E.B. (1992). Crying as social behavior. Infant Mental Health Journal 13, 67–82.

2. Acebo, C., and Thoman, E.B. (1995). Role of infant crying in the early mother-infant dialogue. Physiology and Behavior 57, 541–547.

3. Soltis, J. (2004). The signal functions of early infant crying. Behav Brain Sci. 27, 443–490. 10.1017/s0140525x0400010x.

4. Dudek, J., Faress, A., Bornstein, M.H., and Haley, D.W. (2016). Infant cries rattle adult cognition. PLoS One 11, e0154283. 10.1371/journal.pone.0154283.

5. Sheinkopf, S.J., Iverson, J.M., Rinaldi, M.L., and Lester, B.M. (2012). Atypical cry acoustics in 6-month-old infants at risk for autism spectrum disorder. Autism Res 5, 331–339. 10.1002/aur.1244.

6. Unwin, L.M., Bruz, I., Maybery, M.T., Reynolds, V., Ciccone, N., Dissanayake, C., Hickey, M., and Whitehouse, A.J.O. (2017). Acoustic Properties of Cries in 12-Month Old Infants at High-Risk of Autism Spectrum Disorder. J Autism Dev Disord 47, 2108–2119. 10.1007/s10803-017-3119-z.

7. Esposito, G., and Venuti, P. (2010). Understanding early communication signals in autism: a study of the perception of infants’ cry. J Intellect Disabil Res 54, 216–223. 10.1111/j.1365-2788.2010.01252.x.

8. Kivinummi, A., Naithani, G., Tammela, O., Virtanen, T., Kurkela, E., Alhainen, M., Niehaus, D.J.H., Lachman, A., Lepp&auml;nen, J.M., and Peltola, M.J. (2020). Associations Between Neonatal Cry Acoustics and Visual Attention During the First Year. Front Psychol 11, 577510. 10.3389/fpsyg.2020.577510.

9. LaGasse, L.L., Neal, A.R., and Lester, B.M. (2005). Assessment of infant cry: acoustic cry analysis and parental perception. Mental retardation and developmental disabilities research reviews 11, 83–93. 10.1002/mrdd.20050.

10. Schaefer, G.B., and Mendelsohn, N.J. (2008). Genetics evaluation for the etiologic diagnosis of autism spectrum disorders. Genet Med 10, 4–12. 10.1097/GIM.0b013e31815efdd7.

11. Tordjman, S., Somogyi, E., Coulon, N., Kermarrec, S., Cohen, D., Bronsard, G., Bonnot, O., Weismann-Arcache, C., Botbol, M., Lauth, B., et al. (2014). Gene x environment interactions in autism spectrum disorders: role of epigenetic mechanisms. Frontiers in Psychiatry 5. 10.3389/fpsyt.2014.00053.

12. Moreno-De-Luca, D., and Martin, C.L. (2021). All for one and one for all: heterogeneity of genetic etiologies in neurodevelopmental psychiatric disorders. Curr Opin Genet Dev 68, 71–78. 10.1016/j.gde.2021.02.015.

13. Reichard, J., and Zimmer-Bensch, G. (2021). The Epigenome in Neurodevelopmental Disorders. Front Neurosci 15, 776809. 10.3389/fnins.2021.776809.

14. Scattolin, M.A.A., Resegue, R.M., and Rosário, M.C.D. (2021). The impact of the environment on neurodevelopmental disorders in early childhood. J Pediatr (Rio J). 10.1016/j.jped.2021.11.002.

15. Reichenberg, A., Gross, R., Weiser, M., Bresnahan, M., Silverman, J., Harlap, S., Rabinowitz, J., Shulman, C., Malaspina, D., Lubin, G., et al. (2006). Advancing paternal age and autism. Arch Gen Psychiatry 63(9), 1026–1032. 10.1001/archpsyc.63.9.1026.

16. Hultman, C.M., Sandin, S., Levine, S.Z., Lichtenstein, P., and Reichenberg, A. (2011). Advancing paternal age and risk of autism: new evidence from a population-based study and a meta-analysis of epidemiological studies. Mol Psychiatry 16, 1203–1212. 10.1038/mp.2010.121.

17. Lundstrom, S., Haworth, C.M., Carlstrom, E., Gillberg, C., Mill, J., Rastam, M., Hultman, C.M., Ronald, A., Anckarsater, H., Plomin, R., et al. (2010). Trajectories leading to autism spectrum disorders are affected by paternal age: findings from two nationally representative twin studies. J Child Psychol Psychiatry 51, 850–856. 10.1111/j.1469-7610.2010.02223.x.

18. Hubert, A., Szöke, A., Leboyer, M., and Schürhoff, F. (2011). [Influence of paternal age in schizophrenia]. Encephale 37, 199–206. 10.1016/j.encep.2010.12.005.

19. Kimura, R., Yoshizaki, K., and Osumi, N. (2018). Risk of Neurodevelopmental Disease by Paternal Aging: A Possible Influence of Epigenetic Alteration in Sperm. Adv Exp Med Biol 1012, 75–81. 10.1007/978-981-10-5526-3_8.

20. Krug, A., Wöhr, M., Seffer, D., Rippberger, H., Sungur, A., Dietsche, B., Stein, F., Sivalingam, S., Forstner, A.J., Witt, S.H., et al. (2020). Advanced paternal age as a risk factor for neurodevelopmental disorders: a translational study. Mol Autism 11, 54. 10.1186/s13229-020-00345-2.

21. Yamamoto, T. (2021). Genomic Aberrations Associated with the Pathophysiological Mechanisms of Neurodevelopmental Disorders. Cells 10. 10.3390/cells10092317.

22. Mooney, R. (2020). The neurobiology of innate and learned vocalizations in rodents and songbirds. Current opinion in neurobiology 64, 24–31.

23. von Merten, S., Pfeifle, C., Künzel, S., Hoier, S., and Tautz, D. (2021). A humanized version of Foxp2 affects ultrasonic vocalization in adult female and male mice. Genes Brain Behav 20, e12764. 10.1111/gbb.12764.

24. D’Amato, F., Scalera, E., Sarli, C., and Moles, A. (2005 Jan). Pups call, mothers rush: does maternal responsiveness affect the amount of ultrasonic vocalizations in mouse pups?. Behav Genet. 35 (1), 103–112. 10.1007/s10519-004-0860-9.

25. Hahn, M.E., and Lavooy, M.J. (2005). A Review of the methods of studies on infant ultrasound production and maternal retrieval in small rodents. Behav Genet. 35 (1), 31–52. 10.1007/s10519-004-0854-7.

26. Esposito, G., Hiroi, N., and Scattoni, M.L. (2017). Cry, baby, cry: expression of distress as a biomarker and modulator in autism spectrum disorder. International Journal of Neuropsychopharmacology 20, 498–503. 10.1093/ijnp/pyx014.

27. Grimsley, J.M., Monaghan, J.J., and Wenstrup, J.J. (2011). Development of social vocalizations in mice. PLoS One 6, e17460. 10.1371/journal.pone.0017460.

28. Wohr, M. (2014). Ultrasonic vocalizations in Shank mouse models for autism spectrum disorders: detailed spectrographic analyses and developmental profiles. Neurosci Biobehav Rev 43, 199–212. 10.1016/j.neubiorev.2014.03.021.

29. Dougherty, J.D., Maloney, S.E., Wozniak, D.F., Rieger, M.A., Sonnenblick, L., Coppola, G., Mahieu, N.G., Zhang, J., Cai, J., Patti, G.J., et al. (2013). The disruption of Celf6, a gene identified by translational profiling of serotonergic neurons, results in autism-related behaviors. J Neurosci 33, 2732–2753. 10.1523/JNEUROSCI.4762-12.2013.

30. Shu, W., Cho, J.Y., Jiang, Y., Zhang, M., Weisz, D., Elder, G.A., Schmeidler, J., De Gasperi, R., Sosa, M.A., Rabidou, D., et al. (2005). Altered ultrasonic vocalization in mice with a disruption in the Foxp2 gene. Proc Natl Acad Sci U S A 102, 9643–9648. 10.1073/pnas.0503739102.

31. Scattoni, M.L., Gandhy, S.U., Ricceri, L., and Crawley, J.N. (2008). Unusual repertoire of vocalizations in the BTBR T+tf/J mouse model of autism. PLoS One 3, e3067. 10.1371/journal.pone.0003067.

32. Yoshizaki, K., Furuse, T., Kimura, R., Tucci, V., Kaneda, H., Wakana, S., and Osumi, N. (2016). Paternal Aging Affects Behavior in Pax6 Mutant Mice: A Gene/Environment Interaction in Understanding Neurodevelopmental Disorders. PloS one 11, e0166665. 10.1371/journal.pone.0166665.

33. Yoshizaki, K., Kimura, R., Kobayashi, H., Oki, S., Kikkawa, T., Mai, L., Koike, K., Mochizuki, K., Inada, H., Matsui, Y., et al. (2021). Paternal age affects offspring via an epigenetic mechanism involving REST/NRSF. EMBO Rep 22, e51524. 10.15252/embr.202051524.

34. Tachibana, R.O., Kanno, K., Okabe, S., Kobayasi, K.I., and Okanoya, K. (2020). USVSEG: A robust method for segmentation of ultrasonic vocalizations in rodents. PLoS One 15, e0228907. 10.1371/journal.pone.0228907.

35. Fonseca, A.H., Santana, G.M., Bosque Ortiz, G.M., Bampi, S., and Dietrich, M.O. (2021). Analysis of ultrasonic vocalizations from mice using computer vision and machine learning. Elife 10. 10.7554/eLife.59161.

36. Goffinet, J., Brudner, S., Mooney, R., and Pearson, J. (2021). Low-dimensional learned feature spaces quantify individual and group differences in vocal repertoires. Elife 10. 10.7554/eLife.67855.

37. Hori, K., Yamashiro, K., Nagai, T., Shan, W., Egusa, S.F., Shimaoka, K., Kuniishi, H., Sekiguchi, M., Go, Y., Tatsumoto, S., et al. (2020). AUTS2 Regulation of Synapses for Proper Synaptic Inputs and Social Communication. iScience 23, 101183. 10.1016/j.isci.2020.101183.

38. Branchi, I., Santucci, D., and Alleva, E. (2001). Ultrasonic vocalisation emitted by infant rodents: a tool for assessment of neurobehavioural development. Behav Brain Res 125, 49–56. 10.1016/s0166-4328(01)00277-7.

39. Scattoni, M.L., Crawley, J., and Ricceri, L. (2009). Ultrasonic vocalizations: a tool for behavioural phenotyping of mouse models of neurodevelopmental disorders. Neurosci Biobehav Rev 33, 508–515. 10.1016/j.neubiorev.2008.08.003.

40. Patten, E., Belardi, K., Baranek, G.T., Watson, L.R., Labban, J.D., and Oller, D.K. (2014). Vocal patterns in infants with autism spectrum disorder: canonical babbling status and vocalization frequency. J Autism Dev Disord 44, 2413–2428 10.1007/s10803-014-2047-4.

41. Lopatina, O.L., Furuhara, K., Ishihara, K., Salmina, A.B., and Higashida, H. (2017). Communication Impairment in Ultrasonic Vocal Repertoire during the Suckling Period of Cd157 Knockout Mice: Transient Improvement by Oxytocin. Frontiers in Neuroscience 11. 10.3389/fnins.2017.00266.

42. Takahashi, T., Okabe, S., Broin, P.O., Nishi, A., Ye, K., Beckert, M.V., Izumi, T., Machida, A., Kang, G., Abe, S., et al. (2016). Structure and function of neonatal social communication in a genetic mouse model of autism. Mol Psychiatry 21, 1208–1214. 10.1038/mp.2015.190.

43. Hiramoto, T., Kang, G., Suzuki, G., Satoh, Y., Kucherlapati, R., Watanabe, Y., and Hiroi, N. (2011). Tbx1: identification of a 22q11.2 gene as a risk factor for autism spectrum disorder in a mouse model. Hum Mol Genet 20, 4775–4785. 10.1093/hmg/ddr404.

44. Romano, E., Michetti, C., Caruso, A., Laviola, G., and Scattoni, M.L. (2013). Characterization of neonatal vocal and motor repertoire of reelin mutant mice. PLoS One 8, e64407. 10.1371/journal.pone.0064407.

45. Lai, J.K., Sobala-Drozdowski, M., Zhou, L., Doering, L.C., Faure, P.A., and Foster, J.A. (2014). Temporal and spectral differences in the ultrasonic vocalizations of fragile X knock out mice during postnatal development. Behav Brain Res 259, 119–130. 10.1016/j.bbr.2013.10.049.

46. Miranda, R., Nagapin, F., Bozon, B., Laroche, S., Aubin, T., and Vaillend, C. (2015). Altered social behavior and ultrasonic communication in the dystrophin-deficient mdx mouse model of Duchenne muscular dystrophy. Mol Autism 6, 60. 10.1186/s13229-015-0053-9.

47. Rice, C.E., Naarden Braun, K.V., Kogan, M.D., Smith, C., Kavanagh, L., Strickland, B., Blumberg, S.J., and (CDC), C.f.D.C.a.P. (2014). Screening for developmental delays among young children--National Survey of Children’s Health, United States, 2007. MMWR Suppl 63, 27–35.

48. Boyle, C.A., Decouflé, P., and Yeargin-Allsopp, M. (1994). Prevalence and health impact of developmental disabilities in US children. Pediatrics 93, 399–403.

49. Rosenberg, S.A., Zhang, D., and Robinson, C.C. (2008). Prevalence of developmental delays and participation in early intervention services for young children. Pediatrics 121, e1503–1509. 10.1542/peds.2007-1680.

50. Zablotsky, B., Black, L.I., Maenner, M.J., Schieve, L.A., Danielson, M.L., Bitsko, R.H., Blumberg, S.J., Kogan, M.D., and Boyle, C.A. (2019). Prevalence and Trends of Developmental Disabilities among Children in the United States: 2009-2017. Pediatrics 144. 10.1542/peds.2019-0811.

51. American Psychiatric Association. (2013). Diagnostic and statistical manual of mental disorders: DSM-5 (5th ed.). (American Psychiatric Association). 10.1176/appi.books.9780890425596.

52. Grzadzinski, R., Huerta, M., and Lord, C. (2013). DSM-5 and autism spectrum disorders (ASDs): an opportunity for identifying ASD subtypes. Mol Autism 4 10.1186/2040-2392-4-12.

53. Wozniak, R.H., Leezenbaum, N.B., Northrup, J.B., West, K.L., and Iverson, J.M. (2017). The development of autism spectrum disorders: variability and causal complexity. Wiley Interdiscip Rev Cogn Sci 8, e1426. 10.1002/wcs.1426.

54. Zwaigenbaum, L., Bryson, S., Rogers, T., Roberts, W., Brian, J., and Szatmari, P. (2005). Behavioral manifestations of autism in the first year of life. Int J Dev Neurosci 23, 143–152. 10.1016/j.ijdevneu.2004.05.001.

55. Piven, J., Elison, J.T., and Zylka, M.J. (2017). Toward a conceptual framework for early brain and behavior development in autism. Mol Psychiatry 22, 1385–1394. 10.1038/mp.2017.131.

56. Ozonoff, S., Losif, A.M., Baguio, F., Cook, I.C., Hill, M.M., Hutman, T., Rogers, S.J., Rozga, A., Sangha, S., Sigman, M., et al. (2010). A Prospective study of the emergence of early behavioral signs of autism. J Am Acad Child Adolesc Psychiary 49, 256–266.

57. Astle, D.E., Holmes, J., Kievit, R., and Gathercole, S.E. (2021). Annual Research Review: The transdiagnostic revolution in neurodevelopmental disorders. J Child Psychol Psychiatry. 10.1111/jcpp.13481.

58. Nakamura, M., Ye, K., E Silva, M.B., Yamauchi, T., Hoeppner, D.J., Fayyazuddin, A., Kang, G., Yuda, E.A., Nagashima, M., Enomoto, S., et al. (2021). Computational identification of variables in neonatal vocalizations predictive for postpubertal social behaviors in a mouse model of 16p11.2 deletion. Mol Psychiatry 26, 6578–6588. 10.1038/s41380-021-01089-y.

59. Alio, A.P., Salihu, H.M., McIntosh, C., August, E.M., Weldeselasse, H., Sanchez, E., and Mbah, A.K. (2012). The effect of paternal age on fetal birth outcomes. Am J Mens Health 6, 427–435. 10.1177/1557988312440718.

60. Khandwala, Y.S., Baker, V.L., Shaw, G.M., Stevenson, D.K., Lu, Y., and Eisenberg, M.L. (2018). Association of paternal age with perinatal outcomes between 2007 and 2016 in the United States: population based cohort study. BMJ 363, k4372. 10.1136/bmj.k4372.

61. Skuse, D., Pickles, A., Wolke, D., and Reilly, S. (1994). Postnatal growth and mental development: evidence for a “sensitive period”. J Child Psychol Psychiatry 35, 521–545. 10.1111/j.1469-7610.1994.tb01738.x.

62. Sanefuji, M., Sonoda, Y., Ito, Y., Ogawa, M., Tocan, V., Inoue, H., Ochiai, M., Shimono, M., Suga, R., Senju, A., et al. (2021). Physical growth and neurodevelopment during the first year of life: a cohort study of the Japan Environment and Children’s Study. BMC Pediatr 21, 360. 10.1186/s12887-021-02815-9.

63. Hashimoto, H., Saito, T.R., Furudate, S., and Takahashi, K. (2001). Prolactin levels and maternal behavior induced by ultrasonic vocalizations of the rat pup. Exp. Anim 50, 307–312. 10.1538/expanim.50.307.

64. Liu, R.C., Miller, K.D., Merzenich, M.M., and Schreiner, C.E. (2003). Acoustic variability and distinguishability among mouse ultrasound vocalizations. J Acoust Soc Am 114, 3412–3422. 10.1121/1.1623787.

65. Liu, R.C., Linden, J.F., and Schreiner, C.E. (2006). Improved cortical entrainment to infant communication calls in mothers compared with virgin mice. Eur J Neurosci 23, 3087–3097. 10.1111/j.1460-9568.2006.04840.x.

66. Ehret, G., and Bernecker, C. (1986). Low-frequency sound communication by mouse pups (Mus musculus): wriggling calls release maternal behaviour. Animal Behaviour 34, 821–830. 10.1016/S0003-3472(86)80067-7.

67. Kikusui, T., and Hiroi, N. (2017). A Self-Generated Environmental Factor as a Potential Contributor to Atypical Early Social Communication in Autism. Neuropsychopharmacology 42, 378. 10.1038/npp.2016.225.

68. Mogi, K., Takakuda, A., Tsukamoto, C., Ooyama, R., Okabe, S., Koshida, N., Nagasawa, M., and Kikusui, T. (2017). Mutual mother-infant recognition in mice: The role of pup ultrasonic vocalizations. Behav Brain Res 325, 138–146. 10.1016/j.bbr.2016.08.044.

69. Mogi, K., Nagasawa, M., and Kikusui, T. (2011). Developmental consequences and biological significance of mother-infant bonding. Prog Neuropsychopharmacol Biol Psychiatry 35, 1232–1241. 10.1016/j.pnpbp.2010.08.024.

70. Kato, R., Machida, A., Nomoto, K., Kang, G., Hiramoto, T., Tanigaki, K., Mogi, K., Hiroi, N., and Kikusui, T. (2021). Maternal approach behaviors toward neonatal calls are impaired by mother’s experiences of raising pups with a risk gene variant for autism. Dev Psychobiol 63, 108–113. 10.1002/dev.22006.

71. Centers for Disease Control and Prevention (December 2, 2021). Data & Statistics on Autism Spectrum Disorder. https://www.cdc.gov/ncbddd/autism/data.html.

72. Osumi, N., and Tatehana, M. (2021). Transgenerational epigenetic information through the sperm: Sperm cells not just merely supply half of the genome for new life; they also seem to transmit additional information via epigenetic modifications. EMBO Rep 22, e53539. 10.15252/embr.202153539.

73. Kong, A., Frigge, M.L., Masson, G., Besenbacher, S., Sulem, P., Magnusson, G., Gudjonsson, S.A., Sigurdsson, A., Jonasdottir, A., Wong, W.S., et al. (2012). Rate of de novo mutations and the importance of father’s age to disease risk. Nature 488, 471–475. 10.1038/nature11396.

74. O’Roak, B.J., Vives, L., Girirajan, S., Karakoc, E., Krumm, N., Coe, B.P., Levy, R., Ko, A., Lee, C., Smith, J.D., et al. (2012). Sporadic autism exomes reveal a highly interconnected protein network of de novo mutations. Nature 485, 246–250. 10.1038/nature10989.

75. Flurkey, K., Currer, J., and Harrison, D. (2007). Mouse models in aging research. In The Mouse in Biomedical Research, J. Fox, S. Barthold, M. Davisson, C. Newcomer, F. Quimby, and A. Smith, eds. (Elsevier), pp.637–672.

76. Rieger, M.A., and Dougherty, J.D. (2016). Analysis of within Subjects Variability in Mouse Ultrasonic Vocalization: Pups Exhibit Inconsistent, State-Like Patterns of Call Production. Front Behav Neurosci 10, 182. 10.3389/fnbeh.2016.00182.

77. Yoshizaki, K., Koike, K., Kimura, R., and Osumi, N. (2017). Early postnatal vocalizations predict sociability and spatial memory in C57BL/6J mice: Individual differences in behavioral traits emerge early in development. PLoS One 12, e0186798. 10.1371/journal.pone.0186798.

78. Konishi, S., and Kitagawa, G. (2008). Information Criteria and Statistical Modeling. (Springer-Verlag New York Press).

79. McLachlan, G., and Peel, D. (2000). Finite Mixture Models., 1st Edition (Wiley-Interscience Press).

