## Supplementary Table 1 for "Advanced paternal age diversifies individual trajectories of vocalization patterns in neonatal mice"

The results of mixed model

| Variable | Fixed effect | F value | *p* value |
|---|---|---|---|
| # of distinct syllables | fathers' age | F (1, 13.1) = 9.870 | 0.008 |
| | postnatal day | F (3, 399.2) = 8.733 | < 0.001 |
| | syllable type | F (11, 4532) = 318.096 | < 0.001 |
| | fathers' age × postnatal day | F (3, 399.2) = 2.303 | 0.077 |
| | fathers' age × syllable type | F (11, 4532) = 17.517 | < 0.001 |
| | postnatal day × syllable type | F (33, 4532) = 12.287 | < 0.001 |
| | fathers' age × postnatal day × syllable type | F (33, 4532) = 2.743 | < 0.001 |
| Duration of distinct syllables | fathers' age | F (1, 23.4) = 31.756 | < 0.001 |
| | postnatal day | F (3, 640.6) = 18.974 | < 0.001 |
| | syllable type | F (11, 3020.3) = 658.364 | < 0.001 |
| | fathers' age × postnatal day | F (3, 640.6) = 0.190 | 0.903 |
| | fathers' age × syllable type | F (11, 3020.3) = 4.209 | < 0.001 |
| | postnatal day × syllable type | F (33, 3007.6) = 9.201 | < 0.001 |
| | fathers' age × postnatal day × syllable type | F (33, 3007.6) = 2.518 | < 0.001 |
| Frequency of distinct syllables | fathers' age | F (1, 98.3) = 14.895 | < 0.001 |
| | postnatal day | F (3, 550.9) = 5.108 | 0.002 |
| | syllable type | F (11, 2971.2) = 246.241 | < 0.001 |
| | fathers' age × postnatal day | F (3, 550.9) = 1.108 | 0.345 |
| | fathers' age × syllable type | F (11, 2971.2) = 1.251 | 0.247 |
| | postnatal day × syllable type | F (33, 2955.1) = 17.760 | < 0.001 |
| | fathers' age × postnatal day × syllable type | F (33, 2955.1) = 1.274 | 0.137 |
| Amplitude of distinct syllables | fathers' age | F (1, 18) = 16.598 | < 0.001 |
| | postnatal day | F (3, 591.5) = 106.860 | < 0.001 |
| | syllable type | F (11, 2988.6) = 350.871 | < 0.001 |
| | fathers' age × postnatal day | F (3, 591.5) = 7.220 | < 0.001 |
| | fathers' age × syllable type | F (11, 2988.6) = 2.391 | 0.006 |
| | postnatal day × syllable type | F (33, 2975.8) = 11.394 | < 0.001 |
| | fathers' age × postnatal day × syllable type | F (33, 2975.8) = 2.347 | < 0.001 |
