## Supplementary Table 2 for "Advanced paternal age diversifies individual trajectories of vocalization patterns in neonatal mice"

The AIC number according to the number of cluster.

| | Number of Cluster | | | | |
|---|---|---|---|---|---|
| Parameters | 1 | 2 | 3 | 4 | **5** |
| PC1 + PC2 | 2265.5 | 2229.2 | 2200.5 | 2158.7 | **2132.2** |
| # of calls & duration | 2011.7 | 1937.8 | 1867.9 | 1867.3 | **1796.4** |
| # of syllable types | 1109.8 | 1033.4 | 1024.9 | 1020.1 | **1002.8** |
| Entropy | 1141.3 | 1042.4 | 1009.0 | 997.3 | **966.7** |
