## Supplementary figures and images for "Advanced paternal age diversifies individual trajectories of vocalization patterns in neonatal mice"

### Supplementary Fig. 1

#### Supplementary Figure 1 USVSEG

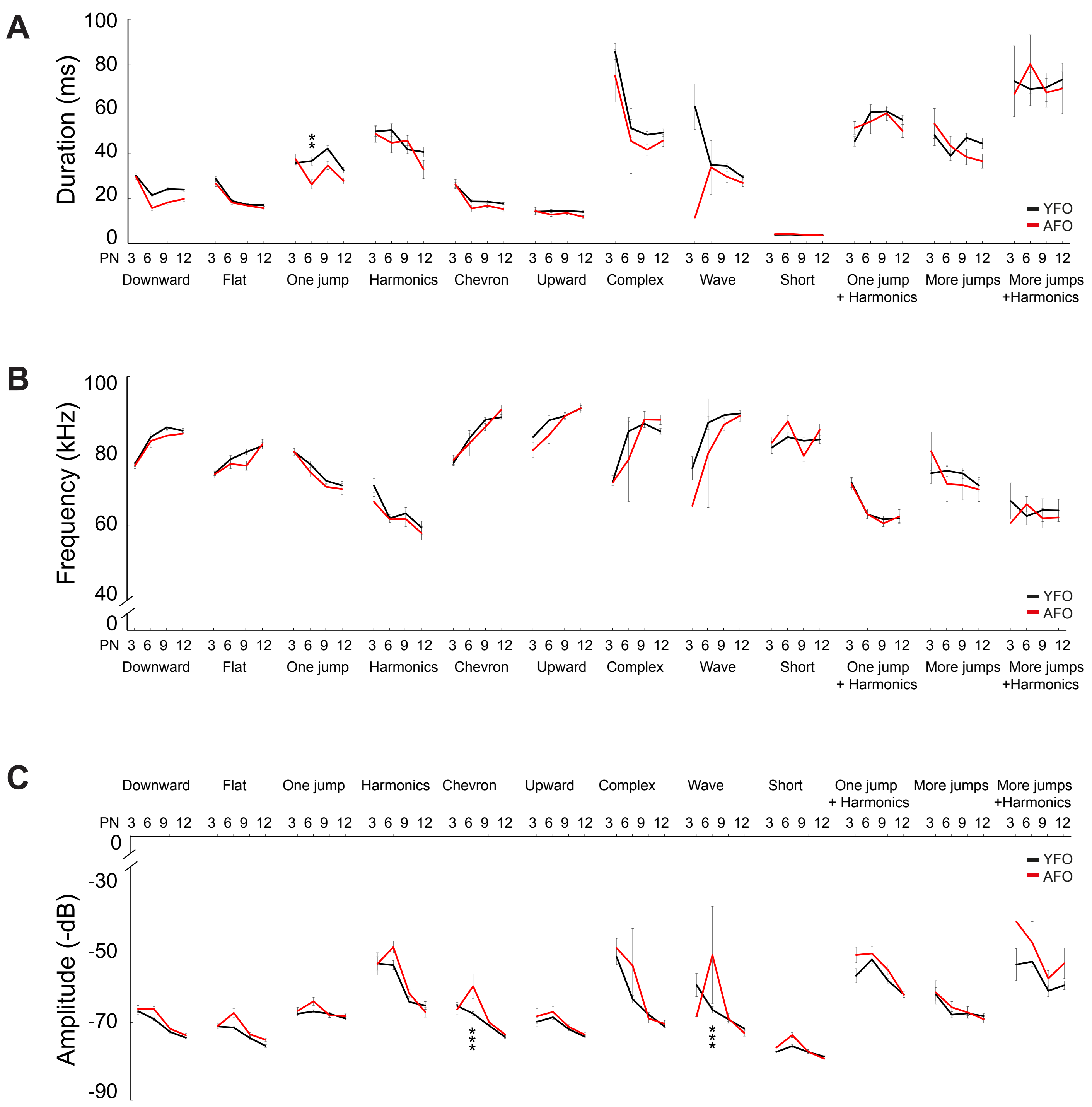

### Supplementary Fig. 2

#### Supplementary Figure 2 USVSEG

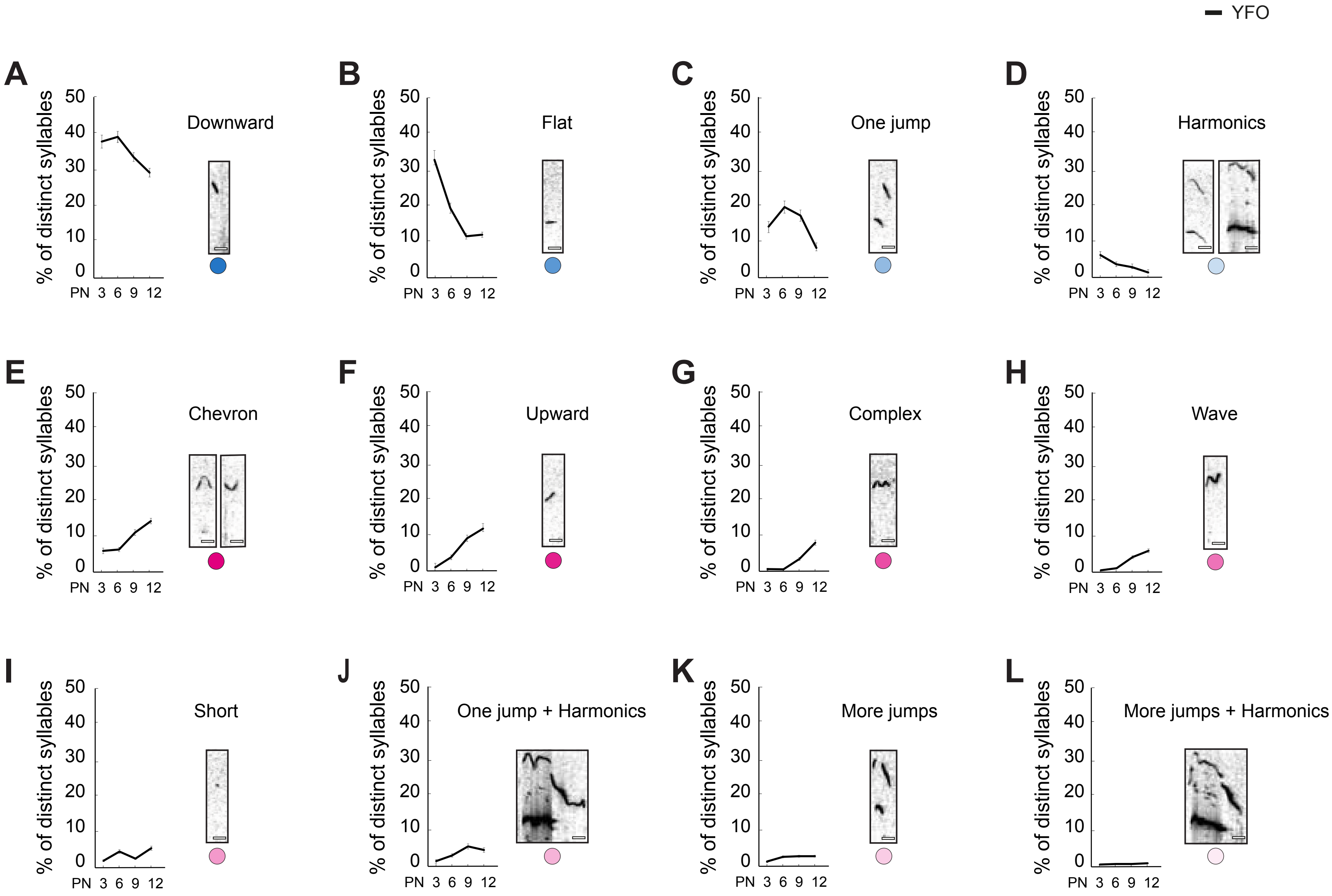

### Supplementary Fig. 3

#### Supplementary Figure 3

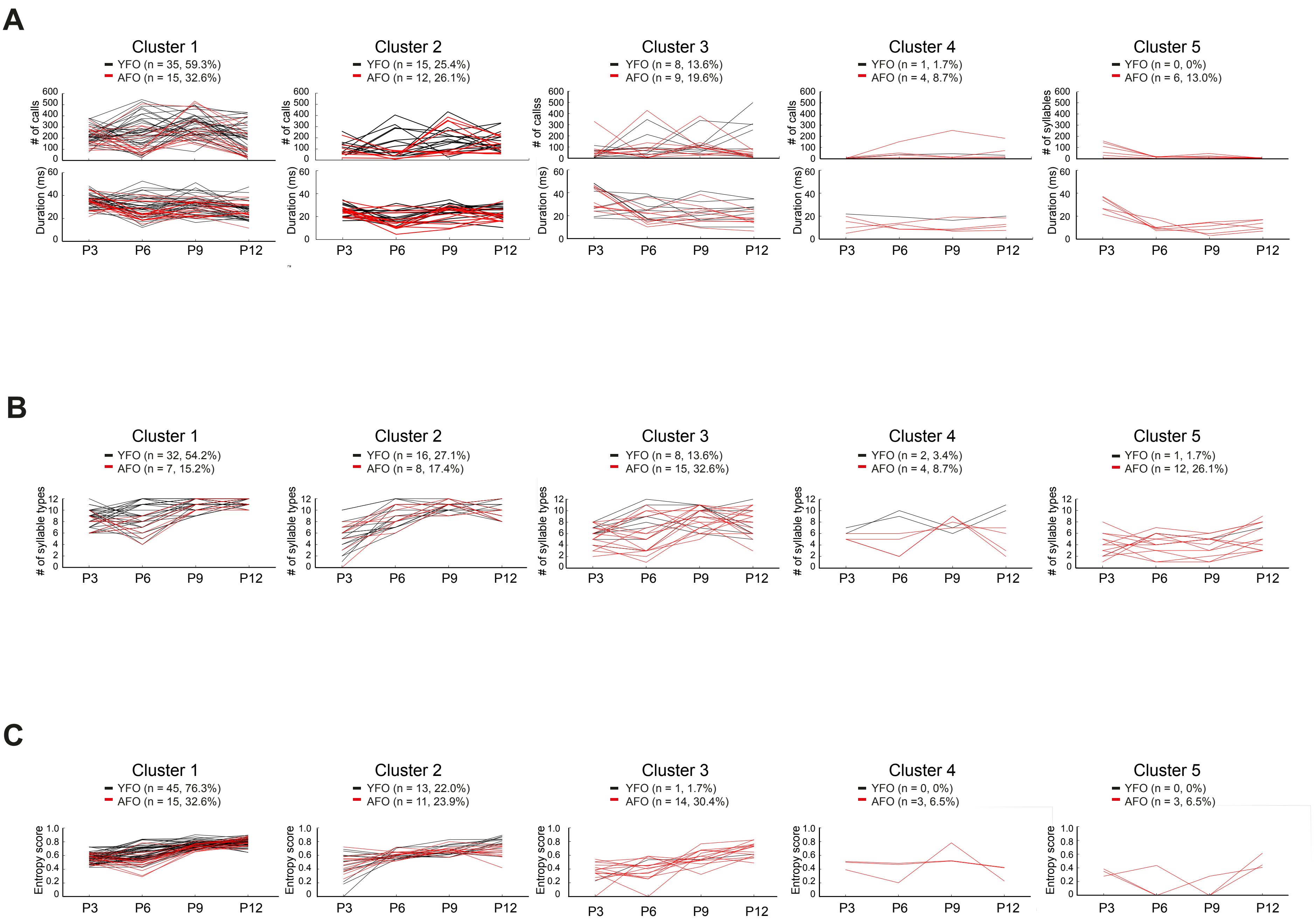

### Supplementary Fig. 4

**Supplementary Figure 4** USVSEG

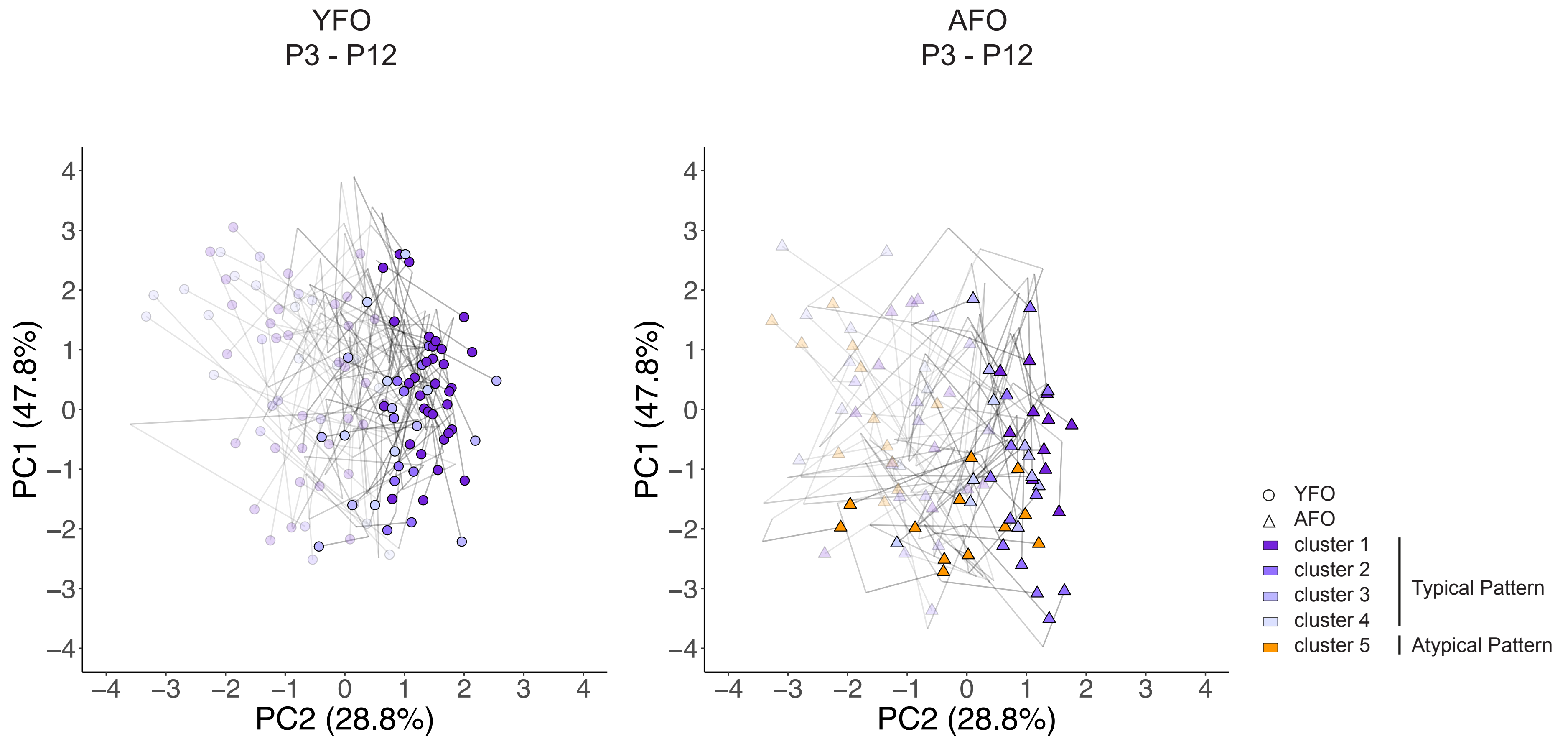

### Supplementary Fig. 5

#### Supplementary Figure 5

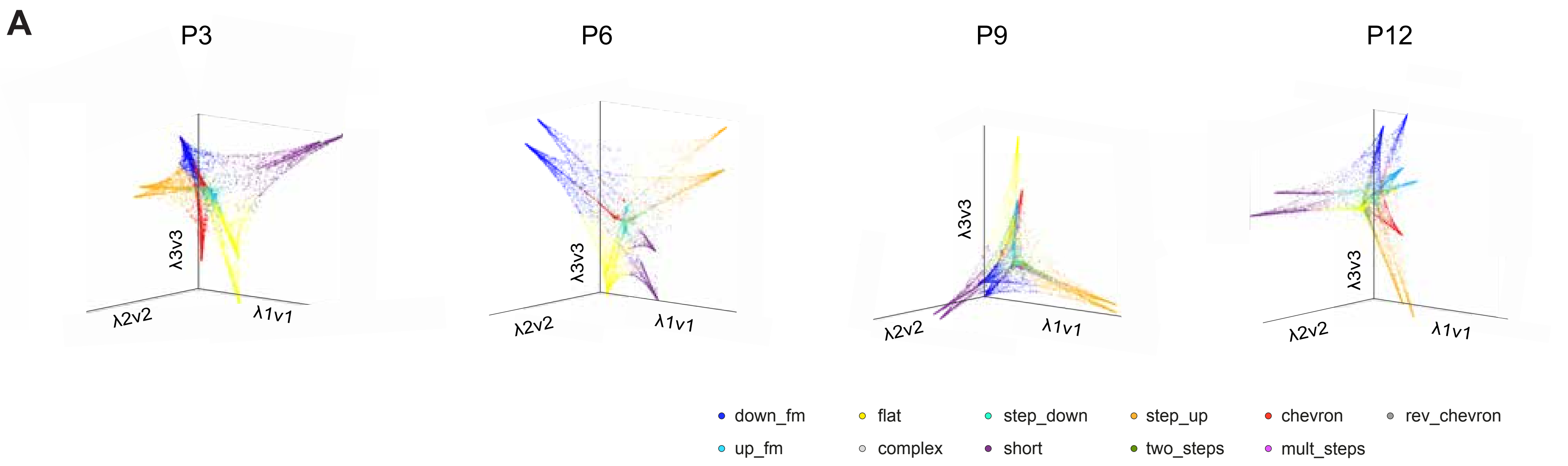


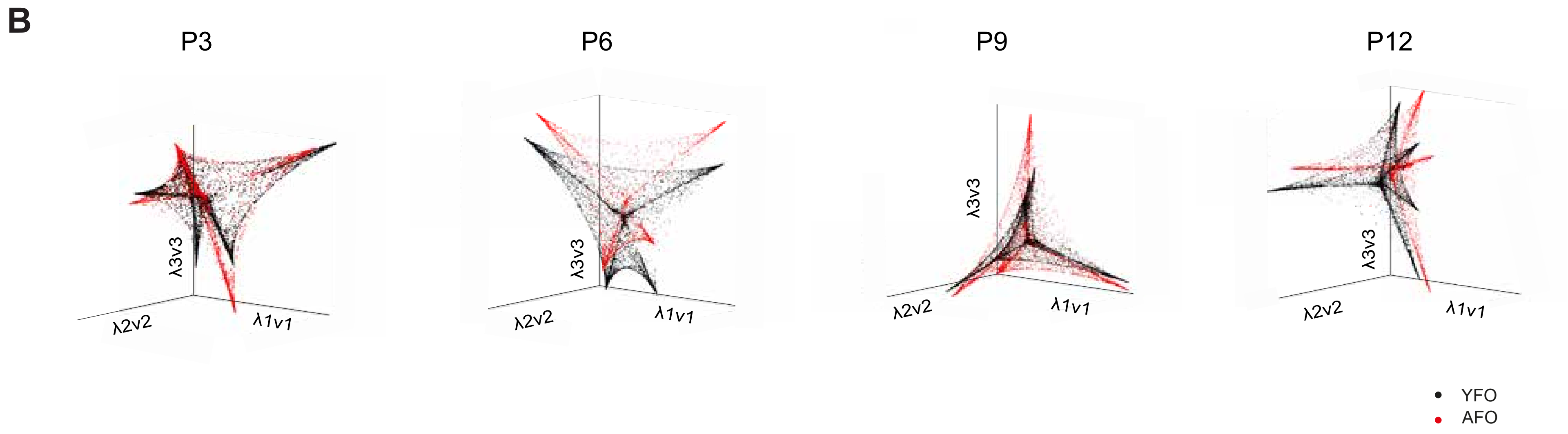
